# Targeting intracranial electrical stimulation to network regions defined within individuals causes network-level effects

**DOI:** 10.1101/2025.07.31.667730

**Authors:** Christopher Cyr, Ania M. Holubecki, Lingxiao Shi, Maya Lakshman, Joseph J. Salvo, Nathan L. Anderson, James E. Kragel, Sarah M. Lurie, Joel Voss, Vasileios Kokkinos, Joshua Rosenow, Stephan U. Schuele, Elizabeth L. Johnson, Christina Zelano, Rodrigo M. Braga

**Author notes:** Corresponding author: Rodrigo M. Braga.

## Abstract

Intracranial electrical stimulation (ES) is routinely used therapeutically, diagnostically, and to provide causal evidence in neuroscience studies. However, our understanding of the brain network-level effects of ES remains limited. We applied precision functional mapping (PFM), based on functional magnetic resonance imaging (fMRI), to define large-scale networks within individual epilepsy patients. We show that single-pulse electrical stimulation (SPES) and high-frequency electrical stimulation (HFES) are more likely to evoke within-network responses and elicit network-related behavioral effects, respectively, when applied near to a PFM-defined network region. Network-level effects were more likely when stimulating sites in white matter, in close proximity to the targeted network, and within a region predominantly occupied by the targeted network. Further, network-specific modulation may be achievable by applying lower current intensities at these sites. Our findings support that modulation of specific networks is achievable by targeting ES to a functional anatomic “sweet spot” that can be identified using PFM.

## Introduction

Intracranial electrical stimulation (ES) is used for therapeutic intervention, diagnosis, neurosurgical planning, and probing brain function (*1–8*). High-frequency electrical stimulation (HFES; ∼50 Hz) can induce or suppress behaviors and is typically used to make inferences about the function of brain regions localized to the stimulation site. In contrast, although single-pulse electrical stimulation (SPES) also affects local brain activity (*9*), it is typically used to make inferences about axonal connections by studying cortico-cortical evoked potentials (CCEPs; *10*–*2*). Thus, HFES and SPES provide useful tools to probe brain function and connectivity, respectively.

The effects of ES are thought to be constrained by the brain’s network architecture (*11*, *13–17*). Thus, prior knowledge of the brain’s network organization, including where specific functions are located and which regions are interconnected, should be useful for guiding ES towards more efficacious modulation of specific networks and functions. This could improve clinical ES mapping as well as inform ES research into brain modulation and function. Functional connectivity (FC), based on resting-state functional magnetic resonance imaging (fMRI; *18*), provides a non-invasive means to map brain networks in humans that can be applied prior to surgical implantation of electrodes. FC-based studies have demonstrated that the human cortex is organized into large-scale networks (*19–21*) which recapitulate axonal projection patterns seen in the macaque (*22–25*) and marmoset (*26*). This is compellingly demonstrated by the observation that disconnection of known polysynaptic cortico-cerebellar circuits following pontine lesions can disrupt FC patterns (*27*). Further, although typically defined from task-free “resting-state” data in humans (i.e., while participants are lying passively in the scanner), networks mapped with FC correspond well with patterns of task-evoked activity, demonstrating that FC can map functional domains (*28*). Thus, FC provides a window into both the regional functional organization of the brain and which regions are interconnected into networks. This suggests that FC-defined brain network maps could provide useful information for targeting intracranial ES to modulate specific functions, as well as predicting the local and distal responses evoked by stimulating a given brain region.

However, this has been difficult to establish, in part because FC estimates have low reliability when defined using limited data per participant (*29*). To overcome this limitation, historically the field has relied on averaging data across individuals to boost reliability; however, group-averaging emphasizes common tendencies and results in maps which do not respect the details of functional organization found within a given individual (*30–32*). This is especially important as functional anatomy can change across the lifespan and patient groups (*33*, *34*). Nonetheless, a correspondence has previously been reported between FC patterns and SPES-evoked potentials (*13–16*), and HFES-elicited behavioral effects (*35*, *17*). These studies show that seed-based or point-to-point FC (e.g., between stimulation site and recording sites) can account for ∼20-40% of the variance in SPES-evoked activity patterns (*13*, *14*). Additionally, interpreting intracranial ES effects with a group-averaged FC-based atlases of brain networks (*21*) supports that SPES-evoked potentials are faster within networks (*15*, *16*), and that HFES effects change when different networks are stimulated (*17*).

Building on these studies, a more precise definition of the location of large-scale network regions within the individual brain should improve our understanding of network-level effects of intracranial ES. This is because a given stimulation site may be defined to be in a given network in a group atlas but could actually be in a completely different network in the stimulated individual due to their idiosyncratic brain organization. Thus, mapping brain networks within individuals would allow assessment of under what functional and anatomical locations stimulation of specific networks may be possible and provide a more accurate representation of the local functional environment that is being perturbed by the applied stimulation.

These considerations motivate a systematic investigation of whether intracranial ES can achieve modulation of specific networks when targeted to networks defined within individuals. Recently, precision functional mapping (PFM), which involves extensive sampling of each individual, has emerged as a means to more reliably and precisely define networks within the individual (*29*, *36–38*). Individualized maps allow a comprehensive assessment of factors affecting the targeting of ES to specific networks. For instance, evidence supports that targeting transcranial magnetic stimulation using individualized FC-defined regions can improve the likelihood of causing therapeutic effects compared to group-based atlases and landmarks (*39*, 40, see also *41*).

Within this context, it is known that the stimulation parameters and anatomical context of the stimulation site also profoundly influence the effects of intracranial ES. Increasing current intensity or pulse width increases the amplitude and spatial extent of evoked responses, and heightens the intensity of perceived behavioral effects (*42–48*). Anatomically, animal and modeling studies of extracellular microstimulation have shown that axons are preferentially affected over cell bodies (*49–56*). Accordingly, intracranial ES applied near or within white matter causes evoked activity patterns with higher amplitude and wider spatial extent than gray matter stimulation (*45*, *47*). This effect is further heightened when stimulation is applied in white matter near the boundary with gray matter (*46*). Additionally, some targeted effects of stimulation are dependent on proximity of the stimulation site to white matter, including network mediated activity (*57*), enhancement of memory specificity (*58*, *8*), and therapeutic benefits for depression (*59*). Thus, when targeting stimulation to network regions defined along the cortical ribbon, the stimulation parameters and anatomical context of the stimulation site are also likely to be important considerations.

Here, we defined large-scale brain networks using PFM in individual participants scheduled to undergo intracranial ES (**Fig. 1A**). Extensive fMRI data were collected from each participant, allowing within-individual network definition using validated procedures (*36*, *60–64*). Following intracranial ES, we assessed how the proximity and dominance of a given network region at the stimulation site influenced evoked effects, and under which stimulation parameters and anatomical contexts network-level effects were more likely.

Our findings reveal that network-level evoked responses and network-related behavioral effects were more likely when stimulation was applied within white matter and close to a PFM-defined network region. Additionally, network-specific effects of SPES were more likely when low current intensity (∼1 mA) was applied at these functional-anatomical locations.

**Fig.1:**
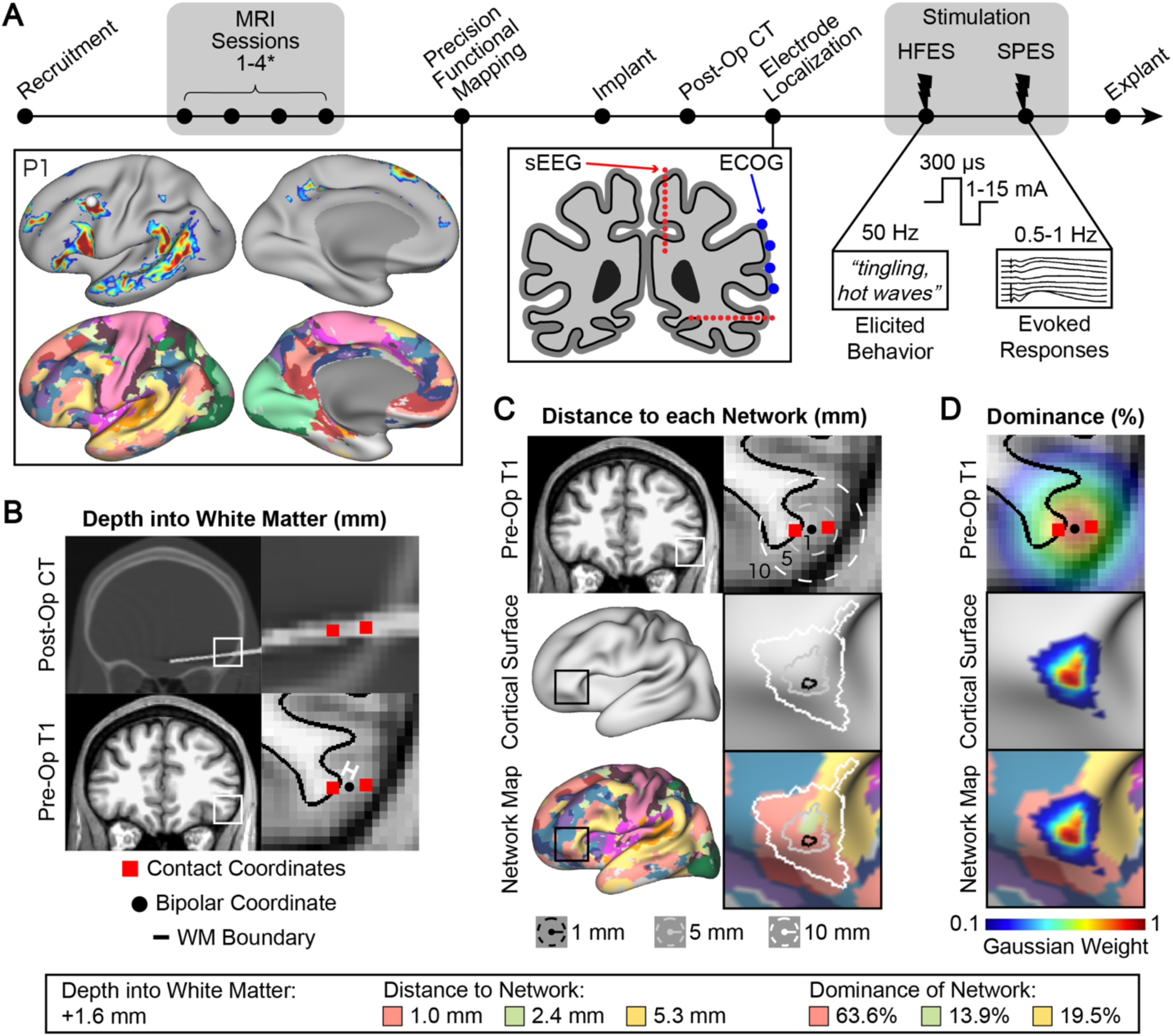
Study overview and derived metrics. (A) Timeline of study. Epilepsy surgery patients were recruited for a precision functional mapping (PFM) study where large-scale networks were defined in each individual prior to intracranial electrical stimulation (ES). Each participant underwent up to four magnetic resonance imaging (MRI) sessions including functional MRI resting-state runs for network definition using functional connectivity (FC). Seed- and parcellation-based maps are shown for example patient P1. Subdural (electrocorticography or ECOG) and depth (stereo electroencephalography or sEEG) electrode locations were chosen strictly for clinical purposes and were determined using a post-operative computed tomography (CT) scan. Intracranial ES was applied at different bipolar stimulation sites, current intensities (1-15mA), and frequencies (SPES: 0.5/1 Hz; HFES: 50 Hz). We conducted a post-hoc analysis of which factors influenced network-level effects of intracranial ES, focusing on (B) the Euclidean distance between the stimulation site (bipolar centroid) and the nearest gray-white matter boundary (defined as negative if in white matter), (C) the shortest (Euclidean) distance from the stimulation site to each PFM-defined network region; and (D) the “Dominance” of each network at the stimulation site, defined by centering a 10-mm full-width at half-maximum (FWHM) Gaussian on the bipolar stimulation site, then calculating the percentage of weighted vertices belonging to each network. We assessed how these stimulation site properties affected network-level evoked responses and behavioral effects of intracranial ES.

Although group atlas and individualized estimates of brain networks were largely overlapped, the HFES sites causing network-related effects were comparatively closer to the analog PFM-defined network than the relevant group atlas-defined network. The results support that non-invasive, individualized definition of large-scale networks allows for network-level targeting of ES.

## Results

### Large-scale networks were successfully defined in presurgical epilepsy patients

Overall, patients provided fMRI data of good tSNR (> 50) across the cortex besides regions near anterior medial temporal lobe (MTL) and ventromedial prefrontal cortex (vmPFC) which typically suffer from signal dropout (**Supp. Fig. S1**). Our data successfully defined large-scale networks within individuals diagnosed with epilepsy (**Fig. 2 & Supp. Fig. S6**), despite the structural abnormalities (dysplasia, sclerosis, etc.) and the characteristically atypical brain activity present in this cohort. These network maps included broad recapitulation of features of the language network (LANG), as well as DN-A and DN-B, which we have extensively studied in prior work on neurotypical controls (e.g., *65*–*67*, *62*–*64*; see also maps of the salience, SAL, and frontoparietal control networks, FPN-A and FPN-B, in *63*, *36*, *61*). However, we also saw idiosyncrasies that are not typically observed in healthy populations that might be of interest to further research. For example, DN-A was missing a medial prefrontal cortex region for P6, P7, and P10 (**Fig. 2 & Supp Fig. S6**). Similarly, we typically expect the LANG network to extend as a strip along the lateral temporal cortex towards the temporopolar region; however, in some subjects this strip was truncated (e.g., see P4 & P10). We circle back to these in the discussion. Despite these observations, individualized network definition was successful and broadly captured expected topographical features of each network.

**Fig 2:**
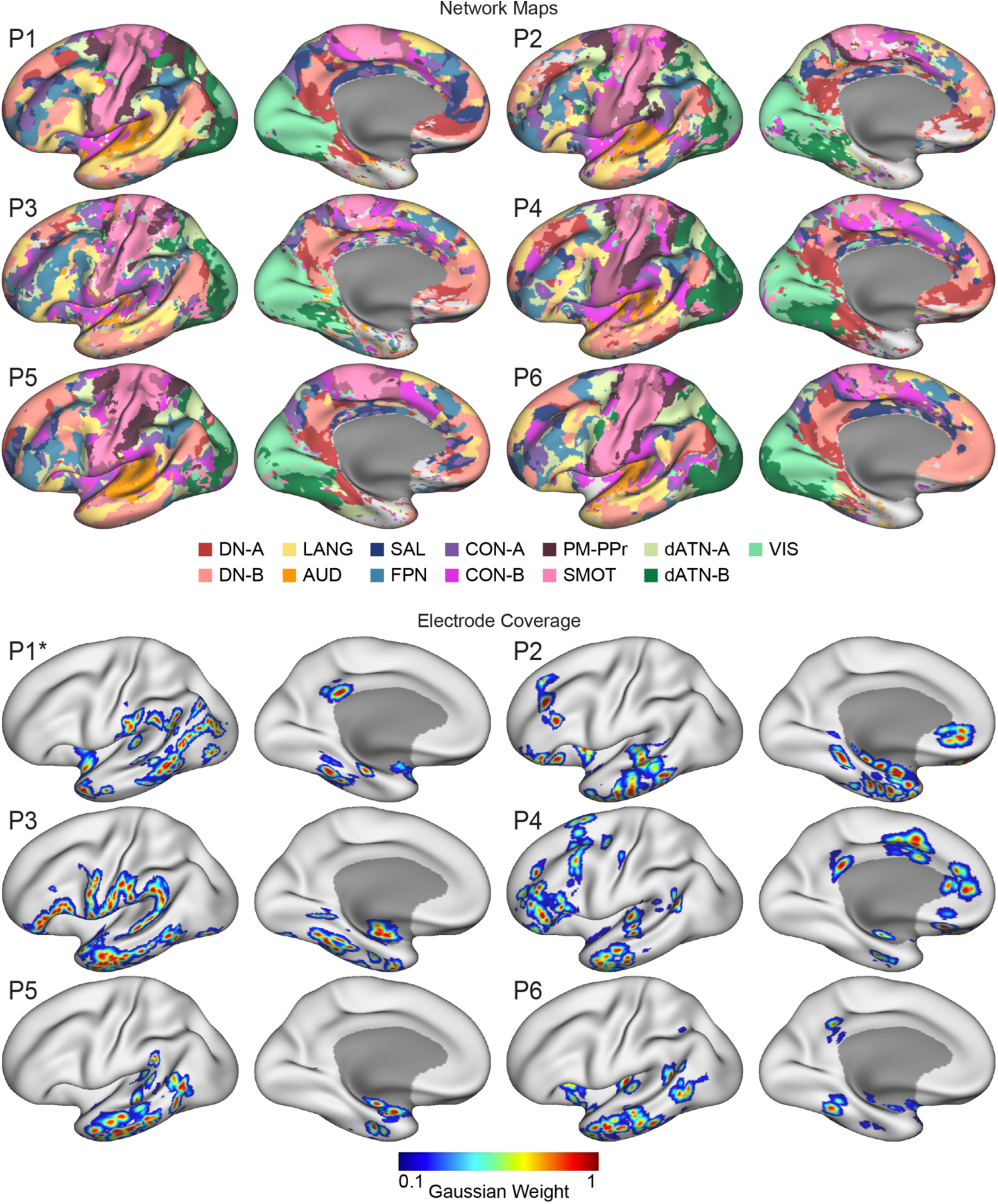
Precision functional mapping estimates of large-scale networks and electrode coverage in 6 example subjects (P1-P6). The remaining 5 subjects, which included right hemisphere and bilateral implants, are shown in Supp. Figs. S6 & S4. Precision functional mapping was used to define large-scale networks in each individual and allowed the localization of electrodes within specific networks. Inflated representations of the left cortical hemisphere are shown in lateral (left) and medial (right) views for a subset of participants (P1 – P6) (Top) Individualized network estimates were derived using data-driven clustering data (*65*) of the extensively collected fMRI data. The initial network maps (shown in Supp. Fig. S2) were then processed post-hoc to remove speckling (small clusters indicating noise) and to combine two networks, somatomotor network A (SMOT-A) and -B (SMOT-B), to homogenize these across individuals. (Bottom) Locations of implanted electrodes are represented using a 10-mm FWHM Gaussian centered on each bipolar contact pair. *Patient P1 had two surgeries which are combined here. DN-A: default network A, DN-B: default network B, LANG: language network, AUD: auditory network, SAL: salience network, FPN: frontoparietal control network, CON-A: cingulo-opercular network A, CON-B: cingulo-opercular network B, SMOT-A: somatomotor network A, SMOT-B: somatomotor network B, PM-PPr: premotor-posterior parietal rostral network, dATN-A: dorsal attention network A, dATN-B: dorsal attention network B, VIS: visual network.

### Single-pulse electric stimulation (SPES) effects from previous studies were replicated

We replicated prior reported features of SPES in our data. This included the inverse relationship between CCEP amplitude and Euclidean distance to the stimulation site (linear model, 𝑦 = 𝑥^−2^, 𝑅^2^ = 0.12, p < 0.001; **Supp. Fig. S5B**; *68*, *69*, *46*, *14*). Also, we replicated the positive correlation between CCEP amplitude and BOLD FC (**Supp. Fig. S5F**; (*13*)) with a majority of participants showing larger CCEPs between stimulation sites and response sites that also displayed larger BOLD correlations, after excluding response sites within 20 mm of the stimulation site (one-sided t tests, t([6, 309]) = [1.76, 6.04], p = [0.046, <0.001], Cohen’s d effect size = [0.21, 0.53]). Additionally, we replicated the finding that stimulation within white matter leads to higher amplitude CCEP responses than gray matter stimulation (**Supp. Fig. S5D**; *46*). Our results did not replicate the findings of (*46*) that local responses are largest when the angle subtended by the stimulation electrode contacts with the cortical columns is around 90° (**Supp. Fig. S5E**). This may be due to the smaller number of stimulation sites included in the current analysis (86 stimulation sites, 8 patients) compared to the (*46*) study (719 stimulation sites, 52 patients). These replications validated that the procedures used here to apply stimulation, localize electrodes, quality control and process the data, and measure evoked responses yielded results that are in line with previous findings, helping to build confidence in the network-based analyses.

### The number of large-scale networks activated by SPES depends on current intensity

The network maps defined in each individual allowed us to assess the factors influencing how many networks are activated by SPES. Recording sites were assigned to the most dominant network in their vicinity using a winner-takes-all method. This process resulted in recordings sites being assigned to a variety of large-scale networks in each participant, demonstrating broad electrode coverage (**Fig. 3A**). After excluding recordings sites without a dominant network (Gaussian-weighted occupancy of at least 10% of this region) and those within 20 mm of the stimulation site, to avoid effects related to volume conduction, we counted the number of networks that included at least one significantly responsive site (trial-averaged maximum amplitude within 20-500 ms post stimulation of Z > 2; **Fig. 3B**).

**Fig. 3:**
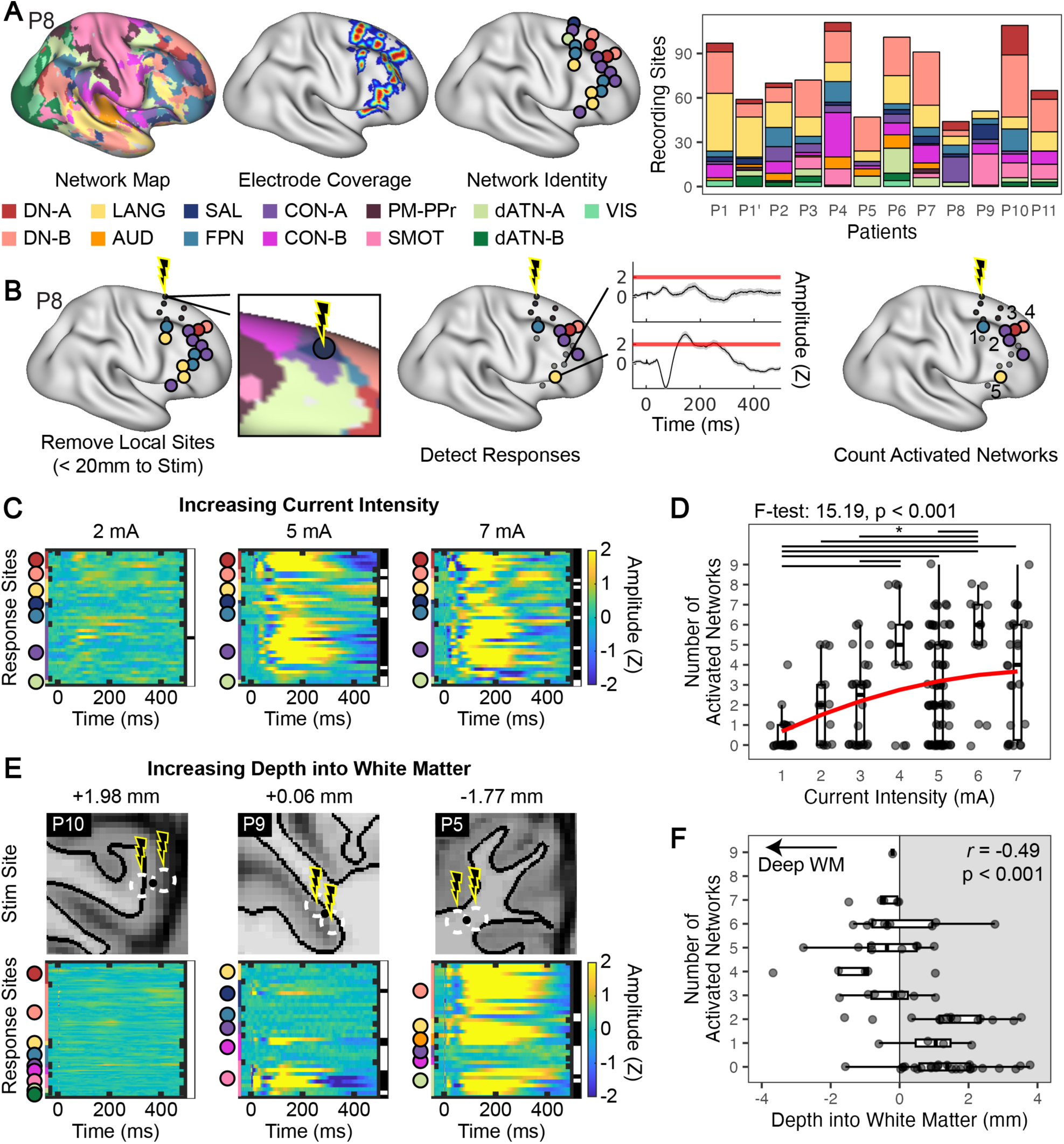
The number of networks activated by SPES depends on the applied current intensity and the depth of the stimulation site into white matter. (A) Recording sites were assigned to the most dominant network in the surrounding region using a winner-takes-all approach. (B) Recording sites within 20 mm (Euclidian) of the stimulation site were excluded, and if at least one distal recording site in a given network showed a significant response (trial-averaged maximum amplitude, 20-500 ms post-stimulation, Z > 2), the network was counted as being activated. (C) Response matrices for distal recording sites (rows) following SPES applied at time 0 ms at the same stimulation site but at three separate current intensities (2, 5, 7 mA). Colored circles and thin bars on the left of each matrix denotes network membership of each recording site. Black stripes on the right denote a significant response. (D) Across stimulation sites, increasing current intensity led to more networks being activated, with a plateau after ≥4 mA. Data were fit to a generalized linear model (red curve) and performance was compared to a constant model (top). A Tukey-Kramer test showed no significant difference (p > 0.05) among higher current intensities, except between 5-6 mA. E) SPES response matrices for 5 mA stimulation at three stimulation sites of varying depths into white matter (mm). (F) Across instances of 5mA stimulation, stimulation sites further into/nearer white matter caused more networks to be activated. Results of a Pearson’s correlation are shown in the top right corner.

The number of networks activated increased with the current intensity applied (generalized linear model, second-order polynomial, identity linkage, F-test: 15.19, p < 0.001; **Fig. 3C-D**). Pairwise comparisons showed that the number of networks activated generally increased with current intensity, and plateaued after ∼4 mA, with no significant differences between levels ≥4 mA besides between 5 mA and 6 mA (Tukey-Kramer test, p < 0.05). These findings align with previous studies showing larger amplitude and more widespread CCEPs with increasing current intensity (*42–44*, *46*, *47*). For most stimulation sites and current intensities applied, the resulting activation was non-specific, including responses in more than one network. However, the fitted model also shows that lower current intensity more often resulted in network-specific activation (i.e., activation of 1 network only).

### More networks are activated when SPES is applied near or within white matter

Focusing on 5 mA stimulation, where we had the most instances of stimulation, we found that stimulation sites further into/nearer white matter activated more networks (Pearson’s correlation, *r* = -0.49, p < 0.001; d.f. = 77; **Fig. 3F**). To test whether this effect was due to the distance of the stimulation site to the gray-white matter boundary specifically, as opposed to the deeper sites also potentially being closer to major white matter tracts, we repeated this analysis after removing stimulation sites more than 15 mm from the surface of the brain (defined using FreeSurfer’s “pial-outer-smoothed” surface; *70*). These results supported that stimulation of peripheral white matter activated more networks than gray matter stimulation (Pearson’s correlation, *r* = -0.45, p < 0.001; d.f. = 63; **Supp. Fig. 7A**).

A potential explanation for this effect is that stimulation sites that are deeper into white matter may be proximal to multiple network regions (i.e., on the opposite side of the gyrus or in adjacent gyri; see anatomical images in **Fig. 3E**). To test this, we assessed whether the number of networks activated could be partially explained by the number of networks located within a 5-mm radius of the stimulation site, and observed no effects (Pearson’s correlation, *r* = -0.02, p = 0.82; d.f. = 77; **Supp. Fig. 7B**).

In contrast, our results supported that a related metric, the Network Dominance (i.e., Gaussian-weighted membership) in the gray matter near the stimulation site, did influence the number of networks activated. As the stimulation site became more dominated by a single network, the number of networks with significant distal responses decreased (Pearson’s correlation, *r* = -0.29, p = 0.008; d.f. = 77; **Supp. Fig. 7C**). The dominance metric weighs networks closer to the bipolar centroid with higher values, implying that the presence of multiple networks close to the stimulation site bipolar centroid led to more networks being activated. Conversely, if a stimulation site was dominated by a single network, fewer networks were activated.

Thus, our results indicate that, aside from current intensity, the number of networks activated was a consequence of the depth of the stimulation site into the peripheral white matter as well as the presence of multiple networks close to the site of stimulation.

### Distal within-network responses are more likely when SPES is applied near the targeted network

We next asked how close a stimulation site needs to be to a network region in order to evoke responses at distal regions of the same network. The Euclidean distance from each stimulation site to each surrounding network region (i.e., the “targeted network”) was calculated, leading to multiple distance values for each stimulation site (i.e., one for each nearby network). For each stimulation site-network region distance, we categorized the pattern of significant distal evoked responses based on whether they were (i) within only the targeted network (“Network *N* Only”, **Fig. 4A**), (ii) within multiple networks including the targeted network (“Includes Network *N*”, **Fig. 4B**), (iii) within one or more networks excluding the targeted network (“Other Network(s)”, **Fig. 4C**), or (iv) not detected (“None”). This was done only for networks in which there were recording sites, after exclusion of sites within 20 mm of the stimulation site. We plotted the results separately for each current intensity applied (**Fig. 4D**).

**Fig. 4:**
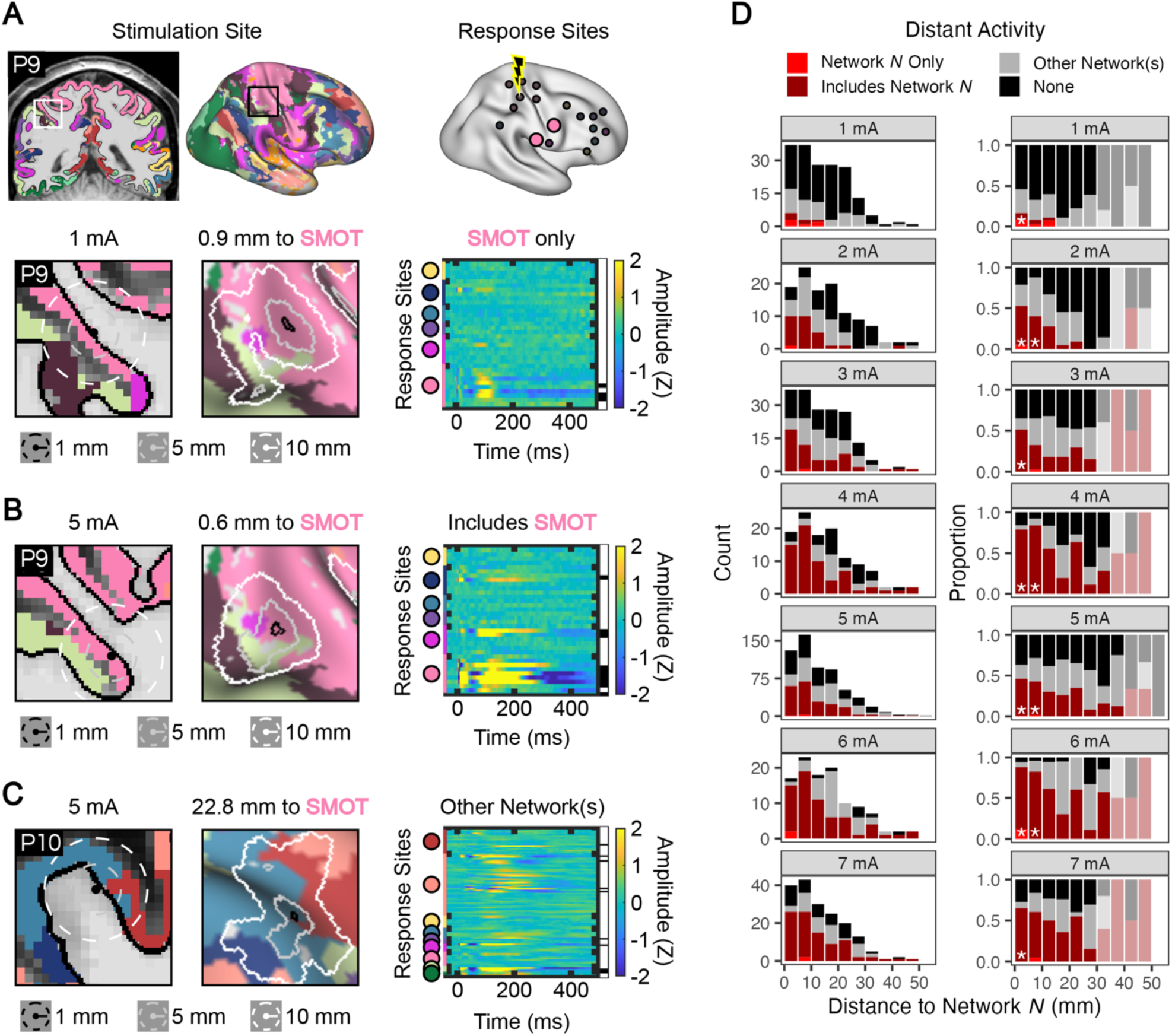
Single-pulse electrical stimulation (SPES) led to distant activation of networks connected to the stimulation site. The Euclidean distance from the stimulation site to a targeted network region was calculated, and the evoked activity pattern was categorized as either showing responses only within the targeted network (A, “Network *N* Only”), within multiple networks including the targeted network (B, “Includes Network *N*”), within one or more networks excluding the targeted network (C, “Other Network(s)”), or not evoking any responses (“None”). This was repeated multiple times per stimulation site for each network in which there were distant recording sites. (D) Count (left column) and proportion (right) of evoked responses in each category binned by distance from the stimulation site to the targeted network. Each plot shows a different applied current intensity. The likelihood of activating a targeted network (red and bright red colors; i.e., Network *N* Only, Includes Network *N*) decreased with increasing distance of the stimulation site to the targeted network. A permutation test showed that only stimulation sites within 10 mm of the targeted network were significantly more likely to cause targeted network effects (i.e., Network *N* Only and Includes Network *N*) than expected by chance (see asterisks on right column, p < 0.05). Current intensity-distance bins with fewer than 5 observations were not assessed (grayed out). Few (16/87) stimulation sites caused network-specific activation (bright red; i.e., Network *N* Only), and these were within short distances (<15 mm) of the targeted network, and more prevalent at low current intensities (∼1 mA).

Note that in these analyses, multiple data points are derived from each stimulation site due to multiple stimulation site-network region distances. For example, a stimulation site that activates one network will lead to one Network *N* only data point as well as several Other Network(s) data points, though each of these data points are assigned to different distance bins depending on the distance to each network. Therefore, we focused on the distribution of response categories detected within each distance bin. The null hypothesis was that distance to the targeted network was not related to activation of the targeted network, meaning that each distance bin should have a similar distribution of response categories.

The likelihood of observing distant responses within a targeted network (i.e., Network *N* Only, plus Includes Network *N*) decreased with increasing distance between the stimulation site and the targeted network (see proportions in **Fig. 4D**, right column). We observed few instances of evoked responses that were specific to the targeted network (Network *N* Only; see bright red in **Fig. 4D**). However, these were more typically observed when the stimulation site was close to the targeted network (<10 mm) and at low current intensity (especially at 1 mA). More often, activation of the targeted network co-occurred with responses in other networks (Includes Network *N*).

To formally test this, we shuffled all the stimulation outcomes (i.e., category labels) within each current intensity to randomize the relationship between stimulation outcome and stimulation site-network region distance. We then recalculated the proportion of each outcome category within the distance bins. This was repeated 10,000 times to create a null distribution of eliciting targeted network effects (i.e., Network *N* Only plus Includes Network *N* categories) at each distance bin and current intensity. If the true likelihood of observing targeted network effects was more extreme than 95% of its respective null distribution, then it was deemed significant. Assessing all current intensity-distance bins with at least 5 observations (i.e., ignoring grayed out columns in Fig. 4D), only stimulation sites that were within 10 mm of a targeted network region were significantly more likely to lead to distant activation of that same network than expected by chance (see asterisks in **Fig. 4D**, right column).

These results support that individualized brain network mapping can predict the distributed spatial pattern of SPES-evoked responses, and that targeting stimulation sites within 10 mm of network regions defined within individuals can lead to distal stimulation of that same network.

### Exploratory analysis of factors necessary for network-specific SPES

Our results supported four factors that might be optimized for achieving modulation of a single network through SPES. First, we found that higher current intensity is more likely to activate more networks (**Fig. 3D**). Second, we found that stimulation near peripheral white matter is particularly efficacious for evoking responses (**Fig. 3F**). Third, we found that stimulation near a functionally defined network region on the gray matter ribbon is more likely to lead to activation of that network (**Fig. 4D**). Fourth, we found that high Network Dominance in the region surrounding the stimulation site is more like to lead to network-specific activation (**Supp. Fig. S7C**). Combining these factors suggests a potential “sweet spot” for achieving network-specific effects, where low current is applied within the white matter, but immediately beneath a dominant (i.e., large) network region mapped within an individual’s cortical ribbon.

We explored whether these factors would allow us to separate the rare instances where we observed network-specific activation from the other stimulation outcomes. We focused on 1 mA stimulation, given the finding that this was more likely to activate one or few networks (**Fig. 3D**) and assessed the remaining parameters. Note that these findings should be seen as exploratory and hypothesis-generating, as our data included few instances of stimulation at 1mA (26 stimulation sites). Statistics are reported for completion, but confirmation of the observed patterns is needed in larger datasets with more instances of low current stimulation (e.g., < 2 mA).

We observed that the stimulation sites causing distal evoked responses (Network *N* Only, Includes Network *N*, or Other Network(s)) all tended to be further into white matter than stimulation sites that did not cause effects (None; **Fig. 5A**), replicating the trend observed at 5 mA (**Fig. 3F**). Pairwise comparisons showed significant differences between the None group and the Includes Network *N* and Other Network(s) groups (Tukey-Kramer test, p < 0.05). Next, we observed that stimulation sites causing effects within the targeted network (Network *N* Only, Includes Network *N*) were all within 15 mm of the targeted network (**Fig. 5B**), while stimulation sites that did not activate the targeted network (Other Network(s), None; **Fig. 5B**) had a wider range of distances. However, pairwise comparisons showed no significant differences between groups (Tukey-Kramer test, p > 0.05).

**Fig. 5:**
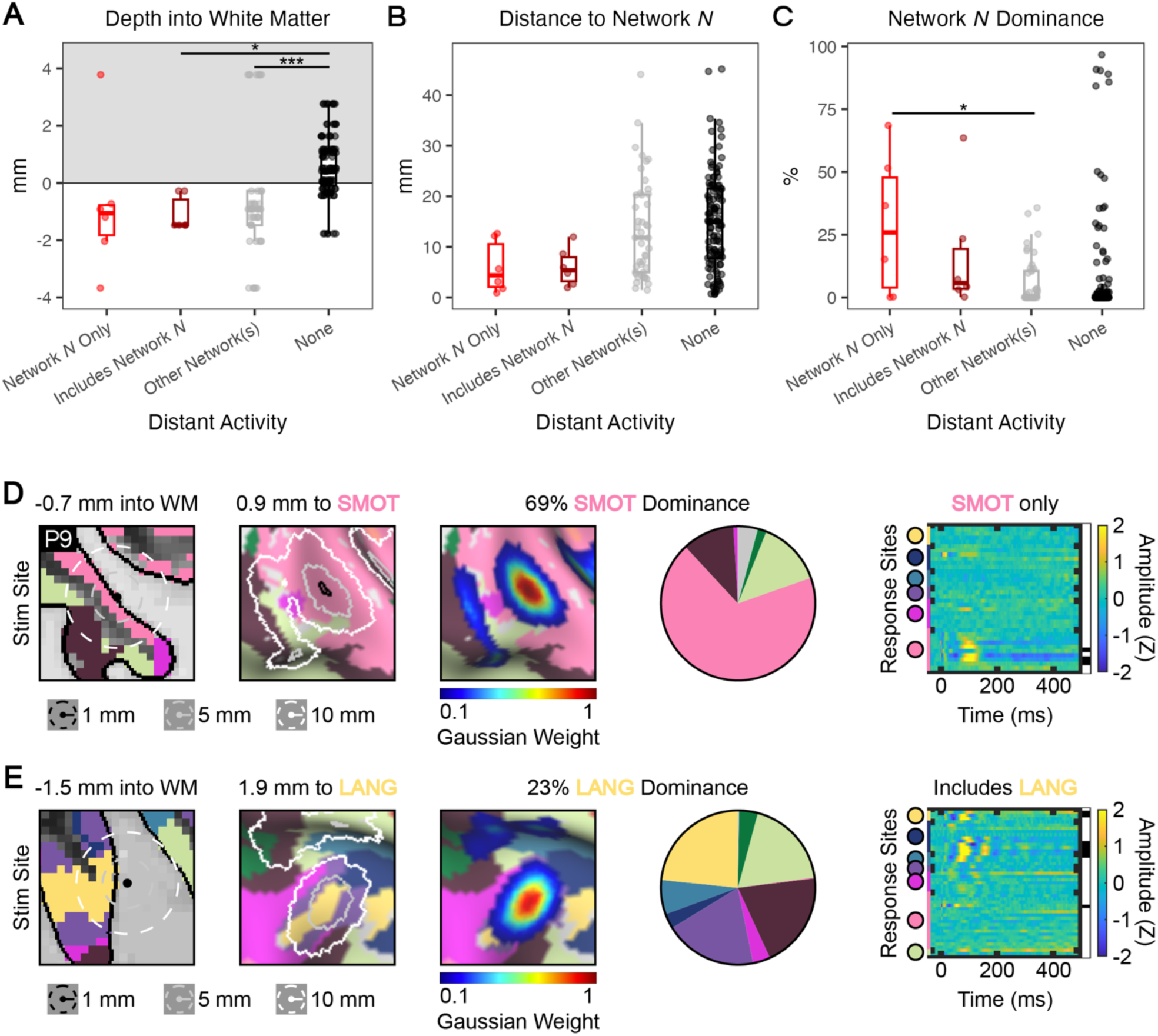
Network-specific responses evoked by low current intensity (1mA) single-pulse electrical stimulation (SPES) are more likely when stimulation is applied near brain regions dominated by the targeted network. Exploratory findings focused on the few instances of low current intensity (1mA) SPES in our dataset revealed that (A) depth into white matter, (B) distance to the targeted network, and (C) dominance of the targeted network at the site of stimulation may be important factors in determining the likelihood of evoking distant activation of only the targeted network (Network *N* Only). Stimulation sites causing targeted network-specific responses (Network *N* Only) tended to be in white matter, near the targeted network, and in brain regions dominated by the targeted network. However, statistics were inconclusive, likely due to the few (26) sites where low current stimulation was applied in our dataset. Pairwise comparisons using a Tukey-Kramer test are shown at the top of each plot (*p < 0.05, ***p < 0.001). The lower panels show example stimulation sites where 1 mA SPES was applied in white matter near regions where (D) one network showed high Network Dominance, leading to network-specific activation of the dominant network, and (E) no network was clearly dominant, leading to nonspecific effects including activation of the targeted network.

Finally, we observed some evidence that distal network-specific activation of the targeted network (Network *N* Only) was observed when stimulation was applied to regions more dominated by the targeted network (**Fig. 5C**). Pairwise comparisons showed a significant difference in targeted Network Dominance between Network *N* Only and Other Network(s) groups (Tukey-Kramer test, p < 0.05) but not other comparisons. Thus, although these results are exploratory, they indicate that achieving targeted network-specific stimulation might benefit from targeting stimulation to the white matter near gray matter regions where a single dominant network region is found (**Fig. 5D**). Conversely, the presence of a balanced distribution of network regions (i.e., low Network Dominance) around the stimulation site may lead to distant activation of multiple networks, even at low current intensity (**Fig. 5E**).

### Depth into white matter affects the likelihood of high-frequency electrical stimulation (HFES) effects

The above results indicated that the current intensity applied and the local environment around the stimulation site, both anatomical (depth into white matter) and functional (proximity to targeted network, dominance of targeted network), are major determinants of downstream effects of SPES. Next, we assessed whether the same principles generalized to HFES (**Fig. 6**).

**Fig. 6:**
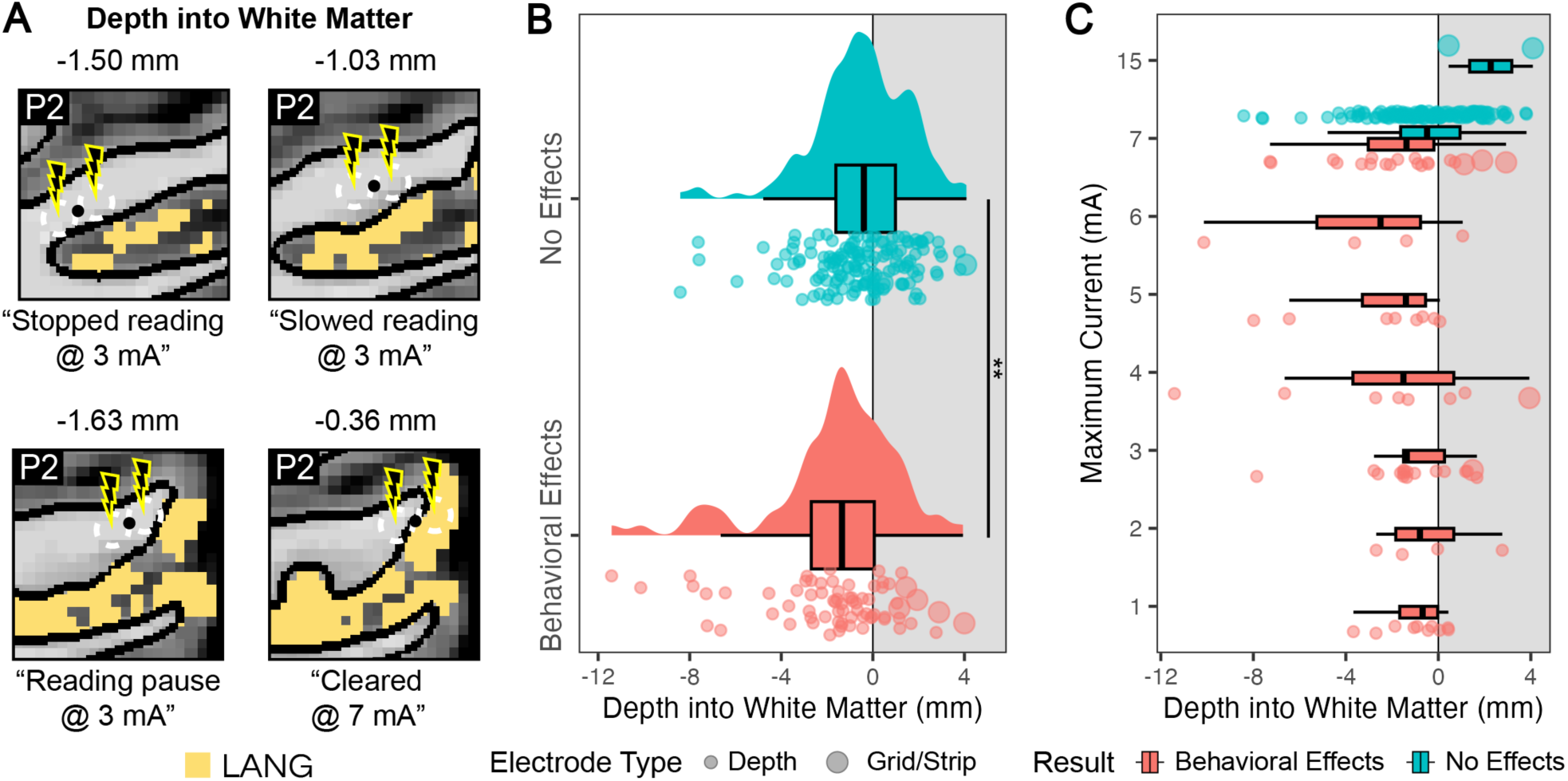
Behavioral effects of high-frequency electrical stimulation (HFES) are influenced by the depth of the stimulation site into white matter. Behavioral effects were categorized as either having elicited behavioral effects (“Behavioral Effects”), or not eliciting effects even at the maximum current intensity that could be applied (“No Effects”, max for depth electrodes: 7mA, max for grid/strip electrodes: 15 mA). Sites that led to seizures, auras, or with behavioral effects concurrent with after discharges were excluded. (A) Examples of HFES performed in participant P2 with language testing near a PFM-defined language network (LANG) region in the lateral temporal cortex (yellow). The site that was closest to gray matter (depth of -0.36 mm) did not elicit effects at 7 mA, while the other sites elicited effects at 3 mA. (B) Across all tested sites, HFES sites that elicited behavioral effects were further into/nearer white matter than those that did not (one-sided t-test ; **p < 0.01). (C) Separating data points by applied current intensity, HFES sites eliciting effects at lower current intensities (∼1mA) appeared to be confined to white matter within 4 mm of the boundary with gray matter, whereas HFES sites not eliciting effects until higher current intensities appeared to be more spread out, though this difference was not statistically significant. Note the higher incidence of gray matter stimulation which elicited no effects despite using higher current intensity.

HFES sites eliciting behavioral effects were significantly further into/nearer white matter than sites that did not elicit effects (one-sided t test, t(106.65) = 3.02, p = 0.002, Cohen’s d effect size = 0.52; **Fig. 6B**). Importantly, this was not due to the current intensity applied for each group. Each site was stimulated with increasing current intensity, and to be included in the No Effects group, stimulation needed to be performed at the highest current intensity allowed (i.e., 7 mA for depth electrodes, 15 mA for grid/strip electrodes; **Fig. 6C**) without eliciting effects. This meant that multiple sites in gray matter failed to elicit behavioral effects even at higher current intensity than was applied to some deeper sites (i.e., where stimulation was stopped at lower current intensity once effects were detected). While there was a significant depth into white matter effect, this pattern was not as clear in the HFES data as it was for SPES. For instance, nearly all of the sites SPES sites that did not cause distant evoked responses at 5 mA (zero networks activated; **Fig. 3F**) were in gray matter. One possible reason for this difference is that some “loss of function” behavioral effects may not have been specifically tested for (e.g., non-linguistic effects) during HFES. Thus, some of the white matter sites in the No Effects group in **Fig. 6B** could have elicited loss of function behavioral effects if tested with a relevant task.

We next assessed the influence of current intensity. Among HFES sites eliciting behavioral effects, those causing effects at lower current intensities (∼1mA) were all confined to within 4 mm of the gray-white matter boundary, whereas those that only elicited behavioral effects at higher current intensities appeared to be more spread out into gray matter and deep white matter (**Fig. 6C**). However, the variances across current intensity levels were not statistically different (Levene’s test, F(6) = 1.52, p = 0.18). In general, the median of the distributions at each current intensity were approximately aligned at a location just below the gray-white matter boundary, consistent with our SPES results and the idea that this location is most efficacious for inducing behavioral effects with HFES.

### Distance to the targeted network region affects behavioral effects of HFES

Our central question was whether the PFM-defined network regions could predict which stimulation sites would cause network-related effects. To test this, we focused on six behavioral effect types (sensorimotor, language, auditory, visual, temperature, and version effects) that were observed as clear effects (i.e. in the absence of other concurrent effects) at more than one stimulation site. We then studied the locations of the stimulation sites causing these effects with respect to specific networks that were hypothesized to be relevant. We expected that stimulation near the SMOT, LANG, and AUD networks, respectively, would elicit sensorimotor, language, and auditory effects. We expected that stimulation near VIS and dATN-B networks, which in combination cover the majority of occipital lobe and the visual streams, would elicit visual effects. Finally, we explored whether other types of behavioral effects were elicited near specific networks, including temperature effects (sensations of warmth, hot flashes, etc.) and version effects (head/eye turning).

**Fig. 7A** shows example participants where the distribution of sites showing clear effects overlapped with the individualized network maps (black outlines). All other subjects and stimulation sites that elicited network-related effects, including the description of effects broken down by category, are shown in **Supp. Figs S8-S15**. Collapsing across networks and associated behavioral domains, the likelihood of observing a behavioral effect related to the targeted network (“Clear Network *N* Effect” plus “Mixed Network *N* Effect”) increased when stimulation was applied closer to the targeted network (**Fig. 7B**). The proportion of these incidences was lower than the SPES results (**Fig. 4D**), though note that some sites may have seemed “silent” because a relevant task was not performed during stimulation. A permutation test confirmed that applying HFES within 5 mm of a targeted network region resulted in above chance likelihood of network-relevant effects.

**Fig. 7:**
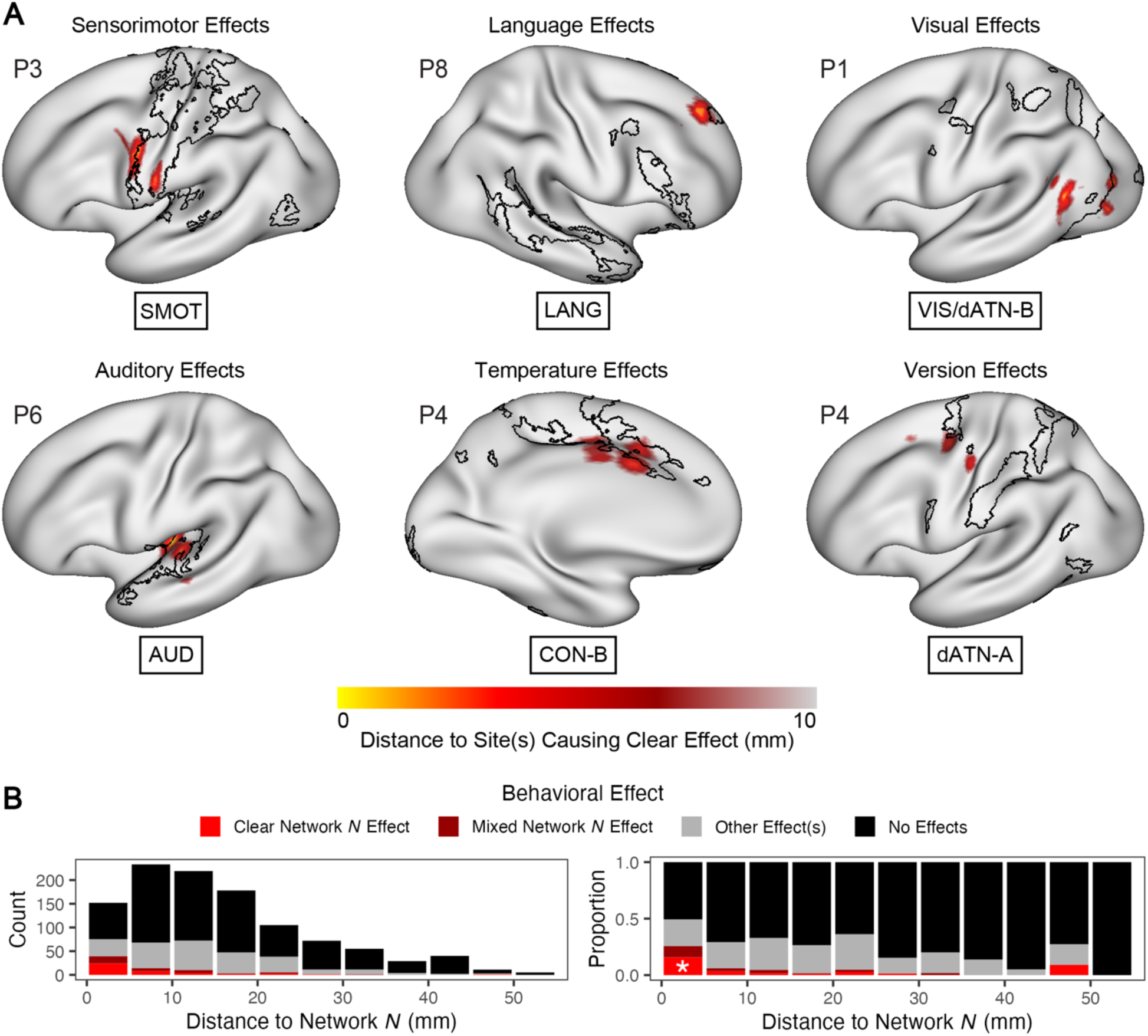
High-frequency electrical stimulation (HFES) elicits behavioral effects more often when applied near behavior-relevant networks. Focusing on HFES sites within 5 mm of gray matter (≥ -5 mm depth into white matter) which are near the functionally mapped networks, HFES effects were assessed in terms of their Euclidean distance to six large-scale networks. (A) Example participants showing the location of sites showing clear network-related effects for each network. Euclidean distance to each stimulation site is shown in the red-yellow colormap, with the relevant network boundaries (SMOT, LANG, etc.) in black. Note that these distance maps can appear discontiguous due to the surface projection step, particularly for contacts that were deeper into the brain. (B) Plot showing the count (left) and proportion (right) of different behavioral effect categories at each distance to the targeted network. Stimulation sites that were closer to the targeted network showed higher incidence network-related behavioral effects (“Clear/Mixed Network *N* Effect”). Only stimulation sites within 5 mm of the targeted network were significantly more likely to cause targeted network effects than expected by chance (see asterisks on right column, derived from a permutation test, p < 0.05).

### Behavioral effects of HFES depend on dominance of the targeted network around the stimulation site

Next, we again assessed whether combining multiple factors would allow us to separate the rare instances of network-specific effects from other HFES outcomes. Dividing the analysis of **Fig. 6B** by outcome, we tested if stimulation sites causing any effects (“Clear Network *N* Effect”, “Mixed Network *N* Effect”, “Other Effect(s)”) tended to be further into white matter than stimulation sites that did not cause effects (“No Effects”; **Fig. 8A**). Pairwise comparisons between categories only showed significant differences between the No Effects group and the Mixed Network *N* Effect and Other Effect(s) groups, but not the Clear Network *N* Effect group (Tukey-Kramer test, p < 0.05; Bonferroni corrected after Fig. 6B analysis).

**Fig. 8:**
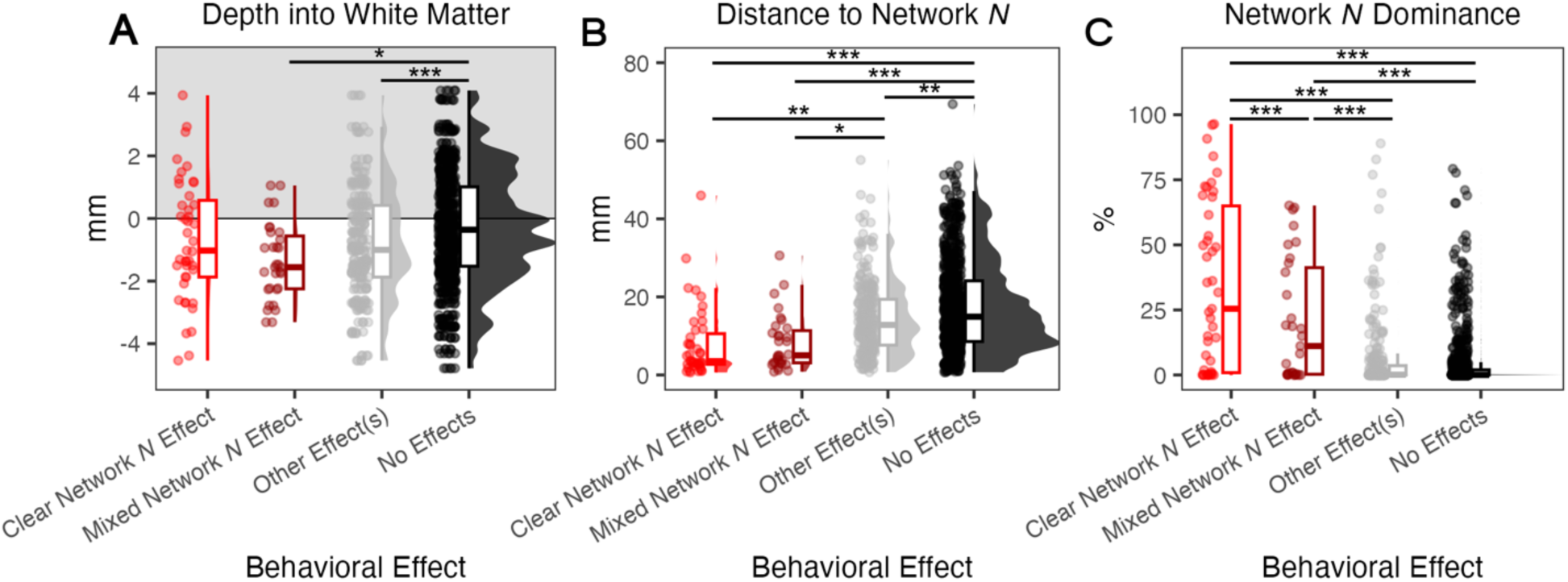
Stimulation sites where high-frequency electrical stimulation (HFES) elicited single-domain behavioral effects were more dominated by the targeted network. Stimulation sites that elicited behavioral effects related to the targeted network (Clear Network *N* Effect, Mixed Network *N* Effect) tended to be located (A) in white matter near the gray matter, and (B) close to the targeted network. Furthermore, (C) stimulation sites that elicited single-domain behavioral effects related to the targeted network (Clear Network *N* Effect) tended to be dominated by the targeted network. This indicates that the functional organization of the stimulated site influences what behavioral effects are elicited. Pairwise comparisons using a Tukey-Kramer test are shown at the top of each plot (*p < 0.05, **p < 0.01, ***p < 0.001).

Next, we observed that stimulation sites causing effects related to the targeted network (Clear Network *N* Effect, Mixed Network *N* Effect) tended to be closer to the targeted network than stimulation sites that did not (Other Effect(s), No Effects; **Fig. 8B**). Pairwise comparisons showed significant differences between all groups besides Clear Network *N* Effect and Mixed Network *N* Effect groups (Tukey-Kramer test, p < 0.05).

Although distance to the targeted network did not differentiate sites that caused clear from mixed effects (**Fig. 8B**), our results showed that stimulation sites causing clear network-related effects (Clear Network *N* Effect) had higher dominance of the relevant targeted network than those eliciting mixed effects (Mixed Network *N* Effect; **Fig. 8C**). Pairwise comparisons showed significant differences between all groups besides Other Effect(s) and No Effects groups (Tukey-Kramer test, p < 0.05). These findings confirmed that the local functional-anatomical architecture at the site of stimulation, including depth into to white matter, distance to specific networks, and dominance of those networks, all influence the effects of SPES and HFES.

### Individualized network maps are more predictive than group-level maps

Precision functional mapping (PFM) requires large amounts of data, raising the question of whether a group-defined atlas can provide sufficient information for targeting intracranial ES without the cost. We assessed whether individualized network estimates derived from PFM provided better predictions of stimulation effects than previous group-level network estimates. To do this, we calculated the distance from each stimulation site to surrounding network regions defined by the 1,000-subject group-level FC atlas from (*21*). We used the 17-network instead of the 7-network atlas because the network sizes in the 17-network atlas were more comparable to those defined within individuals using PFM. For each behavioral effect type, we chose the group-level network analog that was most closely overlapped spatially with the corresponding individualized network (**Fig. 9A**). Note that in some cases, a clear analog is not available. For instance, the closest network to the LANG network in the group atlas is “Default Network B”, while the closest to the AUD network in the group atlas is “Somatomotor B” which includes the ventral somatomotor cortex.

**Fig. 9:**
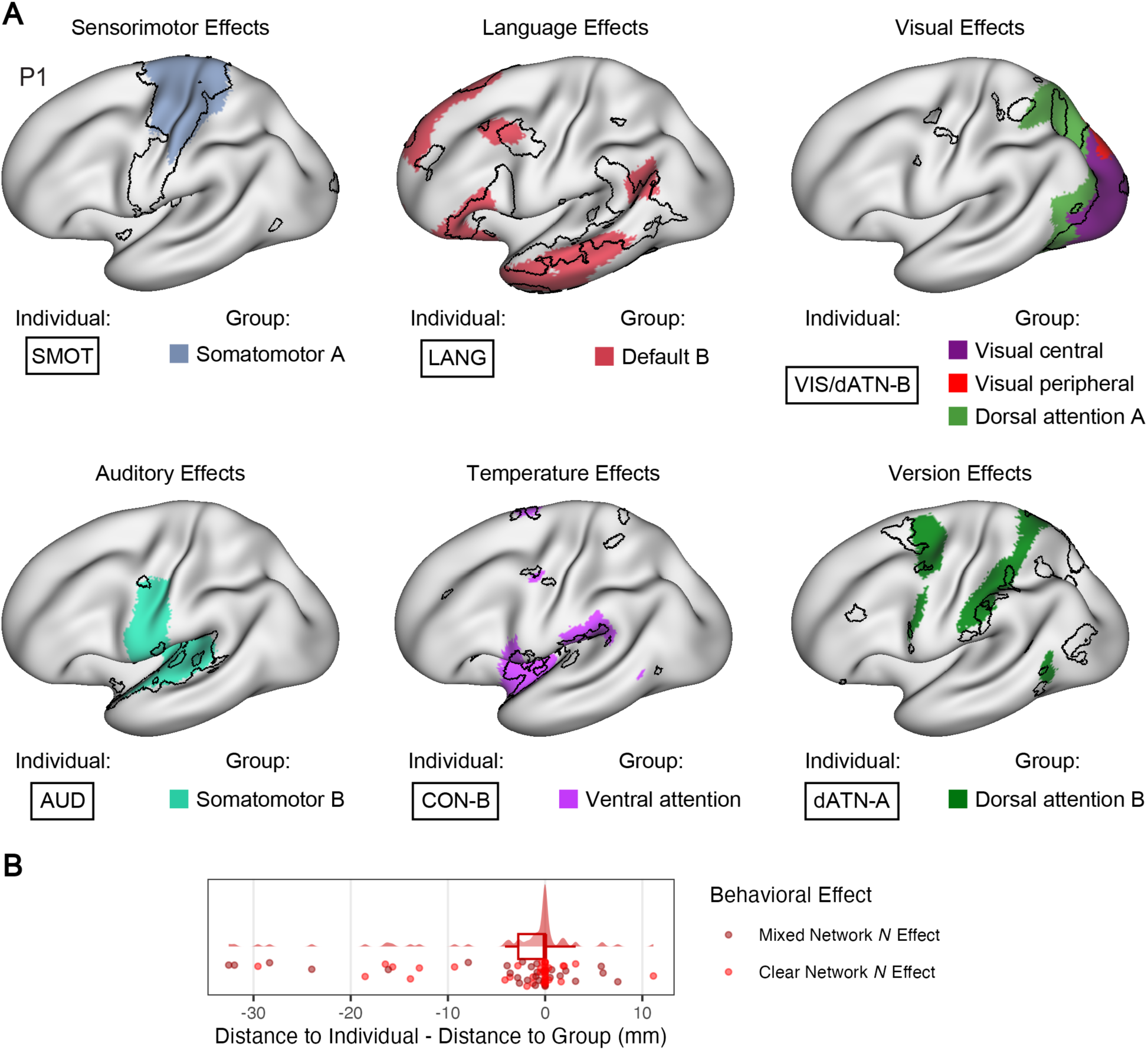
Stimulation sites where high-frequency electrical stimulation (HFES) elicited network-related behavioral effects were closer to the relevant individualized network than a group-level network analog. (A) For each behavioral effect type, group-level network analogs were chosen for each individualized network based on spatial overlap. Group-level network analogs are labeled by their name and color from (*21*). Boundaries of the individualized network are in black. (B) Stimulation sites where HFES elicited network-related behavioral effects were then assessed based on whether they were closer to the relevant individualized network (negative difference in distance) or the group-level network analog (positive difference in distance). The maps from both group and individualized approaches overlapped at most sites (difference in distance equal to 0 mm). However, sites that elicited network-related behavioral effects overall tended to be closer to the individualized networks (i.e., the distribution was shifted towards negative values).

We plotted sites showing a network-related behavioral effect (Clear Network *N* Effect, Mixed Network *N* Effect) against the difference in the distance from the stimulation site to the networks derived from PFM and the group atlas (i.e., distance to individualized PFM network minus distance to group atlas). Sites which overlapped both networks had a difference in distance of 0 mm, and sites closer to the PFM-defined network had a negative difference in distance value. Although many stimulation sites eliciting network-related effects were equally close to both individualized and group-level networks (zero on x-axis), reflecting substantial overlap between the networks, the distribution was negatively shifted (one-sided Wilcoxon signed-rank test, V = 354, p = 0.001; **Fig. 9B**). This supports that stimulation sites overall were closer to the individualized networks, and that the individualized maps more closely captured the sites leading to HFES effects than the group maps.

## Discussion

We mapped large-scale networks within individual participants’ brains using precision functional mapping (PFM) and investigated whether these maps could be used to target intracranial electrical stimulation (ES) to cause network-level evoked responses and network-related behavioral effects (**Fig. 1**). We found that stimulation applied near a targeted network region was more likely to lead to distant activity within other regions of that network (**Figs. 4-5**) and was more likely to elicit behavioral effects associated with that network (**Figs. 7-8**). We further showed that the topography of networks at the stimulation site is also an important factor: activation of distant network regions and clear network-related behavioral effects were more likely when stimulation was applied to brain regions where one network was dominant (**Figs. 5 & 8**). Lastly, we found that PFM was additionally informative in predicting network-related behavioral effects compared to a group-defined network atlas (**Figs. 9**). Thus, our results reveal that PFM can be used to target ES for tailored effects, providing a non-invasive and task-free means to improve the efficacy and accuracy of ES in clinical and research contexts. Based on these results, we outline a proposed framework for achieving network-specific effects through carefully optimized stimulation parameters and targeting of stimulation to specific functional-anatomical sites (**Fig. 10**).

**Fig. 10:**
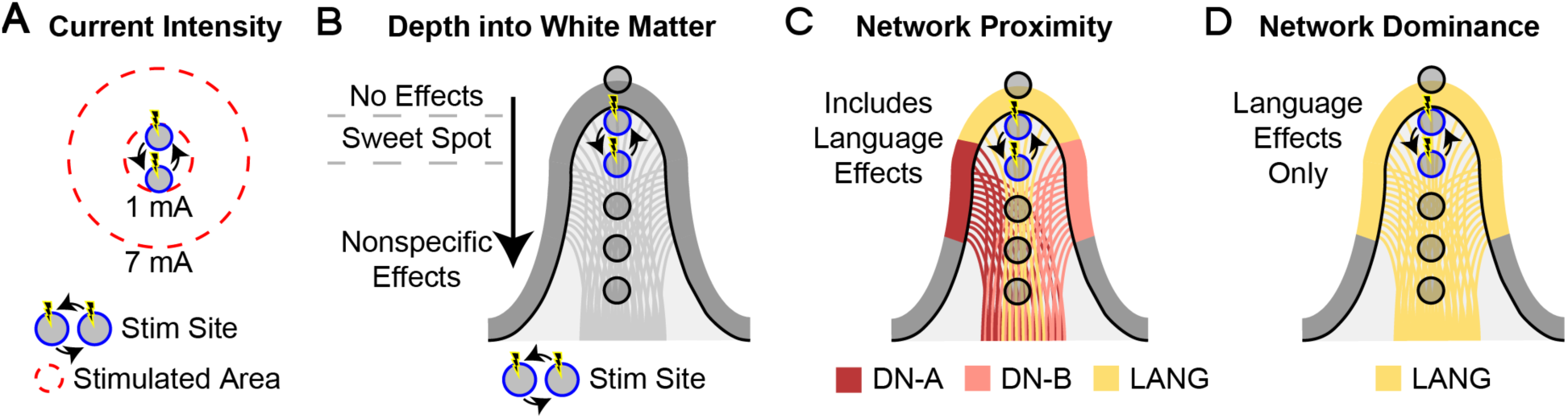
Proposed framework for network-targeted intracranial electrical stimulation. (A) Apply stimulation using low current intensities to reduce the volume of stimulated tissue and increase the likelihood of modulating a single network. (B) Stimulation should be applied within white matter but immediately near (e.g., < 5 mm away) to the gray matter ribbon (i.e., the “sweet spot”) to increase the likelihood of eliciting effects, possibly due to activation of axons leaving the nearby gray matter region. (C) Apply stimulation near a region of the targeted network to increase the likelihood of activating that network and/or eliciting behavioral effects associated with that network. (D) Apply stimulation in a region that is more dominated by the targeted network to increase the likelihood of achieving network-specific activation and effects.

### Precision functional mapping is achievable in individuals with epilepsy

The present results demonstrate that PFM is feasible in individuals with epilepsy, who were able to provide sufficient fMRI data for reliable individualized mapping, in some cases attending multiple MRI sessions. We previously published a case report where PFM was performed in an epilepsy patient prior to invasive monitoring (*71*). Here, we extend this by reporting results from 11 participants who successfully provided between 35 (35) and 189 (147) minutes of collected (retained) fMRI data per patient (**Supp. Table 1**). With recent improvements in fMRI sequences and methods, this approach may be feasible in a wider range of patients.

We found that the final network maps in each participant (**Fig. 2 & Supp. Fig. S6**) preserved anatomical features commonly seen in non-epileptic control subjects (e.g., *36*, *61*, *67*), and which were used here to identify the networks, despite the structural abnormalities and pathological brain activity characteristic of this cohort. This indicates that the epileptic brain is not fundamentally different in its large-scale network organization from the healthy brain, and that PFM can be used to capture individual-specific and age-specific details in cortical organization in epilepsy patients (*30–32*, *34*).

Additionally, because PFM-defined network estimates are congruent between epilepsy patients and healthy controls, recent fMRI findings regarding task-related activations within these networks can inform the selection of specific tasks to administer during HFES. The use of such tasks in a clinical mapping context could enable a higher likelihood of observing loss of function effects based on the stimulation of a given network (*72*, *61*, *67*, *63*, *64*).

However, open questions remain regarding if and how epileptiform activity, which often involves large-amplitude fluctuations of brain activity, alters the organization of individualized network estimates which are defined using similarly slow fluctuations of brain activity (e.g., see third figure of *73*). If so, seizure activity could lead to systematic differences in network topography (*74–76*, *73*, *77*, *78*), particularly during epileptic episodes, but that could persist in interictal periods. Here, none of the participants reported having seizures during the MRI sessions, though it is likely that some epileptiform activity occurred.

We observed some participants who displayed deviations from the expected network organization in certain regions. For instance, in the vmPFC in participants P6, P7, and P10, we did not see an expected region of DN-A in rostral vmPFC (compare with remaining subjects in **Fig. 2 & Supp. Fig. S6;** or to *36*). The vmPFC is densely connected to the anterior MTL and amygdala (*79–82*), which in these patients was their clinically determined site of epileptogenicity (**Supp. Table 1**).

Prior work supports that MTL epilepsy can lead to hypoconnectivity in resting-state networks, particularly the default network which is connected to the MTL (*74*, *83*, *84*, *76–78*). For instance, (*78*) showed that the mPFC in particular displayed decreased functional connectivity in epilepsy patients compared to control participants (with the posteromedial cortex showing the opposite effect). These decreases in functional connectivity, typically reported in group-wise analyses, may appear as topographical differences in the individualized network estimates, including the entire omission of specific network regions as observed here in a few participants. While PFM has shown differences in network topography within brain regions affected by depression (*85*), an open question regards whether PFM can be used to delineate the seizure onset zones based on such topographical quirks (*86*). Future work in this area is warranted.

### Stimulation parameters influence the number of networks activated by SPES

We found that SPES with higher current intensity led to more widespread activation that included responses in many networks (**Fig. 3C & 3D**). This is consistent with prior SPES studies showing increases and subsequent plateauing of responses at higher intensities (≥4 mA; *43*, *44*, *46*). Supporting evidence from animal studies has demonstrated that the current required to activate a neuronal element is proportional to the squared distance from the stimulating electrode (*87–90*). Thus, the radius of neural activation around the stimulation site may expand proportional to current intensity. As this radius expands, stimulation may impact an increasing number of networks that have regions near the stimulation site, leading to propagation of signals along multiple networks (**Fig. 10**). This suggests that network-specific stimulation effects may be more feasible at lower current intensities (e.g., ∼1 mA and below). As shown in **Fig. 3D**, we found that more networks were activated at higher current intensity. Although we had relatively few instances of 1 mA stimulation, the trend in our data supports that lower current intensity was more likely to lead to the network-specific activation.

A second observation was that stimulation applied within white matter tended to activate more networks. Previous studies have shown that the spread of evoked activity increases with increasing depth of the stimulation site into white matter, for both SPES and HFES (*45*, *46*). Other studies have found that the stimulation site must be in or near white matter to elicit intended effects (*57–59*). Supporting evidence from animal and modeling studies of extracellular microstimulation have found that axons are preferentially affected over cell bodies, including at or near the axon initial segment that leaves the cell body (*49–56*). In humans, the strength of interregional evoked responses corresponds to the strength of anatomical connections estimated using diffusion tractography (*10*, *14*, *91*, *92*).

These studies support that stimulation of white matter is particularly efficacious, which raises a conundrum of how to target functional regions that are situated along the cortical ribbon. At the macroscale level, our data suggests that instead of applying stimulation directly to the cortex, moving the stimulation site just below the gray-white matter interface into the white matter may increase the likelihood of eliciting evoked responses and effects. In other words, there may be a sweet spot immediately beneath the gray matter ribbon where stimulation is most likely to cause effects related to functional regions in the adjacent cortex (**Fig. 10B**). This is consistent with our findings.

### Targeting stimulation to network regions increases the likelihood of network-specific effects

We observed that the location of the stimulation site with respect to an individual’s functional topography of large-scale networks could explain stimulation effects. Prior studies have demonstrated that fMRI functional connectivity (FC) corresponds with the spread of effects of ES, both in terms of distal activity evoked by SPES, and behavioral effects elicited by HFES (*13*, *14*, *17*). Here, we were able to leverage PFM to define networks stably within individuals which allowed the localization of electrodes with respect to an individual’s idiosyncratic topography of network regions (**Fig. 2** & **Fig. 3A**). Importantly, this allowed the estimation of the distance between each stimulation or recording site and the nearest region of multiple networks.

We found that SPES was more likely to evoke distant activity in a given network if that network included a region near the stimulation site (**Fig. 4**). Further, when we restricted analyses to the lowest current intensity (1 mA), exploratory results suggested that stimulation was more likely to elicit network-specific responses when the gray matter surrounding the stimulation site was dominated by the targeted network (**Fig. 5**). These findings were based on relatively few datapoints and thus need validation in larger cohorts. However, they were corroborated by results from HFES, where both the proximity and dominance of the targeted network at the stimulation site differentiated instances where network-specific behavioral effects were elicited (i.e., a Clear Network *N* Effect; **Fig. 8**).

Lastly, we directly contrasted the individualized network estimates with group-level estimates, which revealed that the individualized estimates more often captured sites that showed network-related effects of HFES (**Fig. 9**). Thus, our data demonstrate that PFM can be used to target electrical stimulation to specific networks within individuals. Further, our results suggest that precise tailoring of stimulation parameters and the location of stimulation may lead to network-specific modulation. We further emphasize that even though our functional mapping relied on resting-state FC, these network estimates predicted not only the spread of evoked responses but also the likelihood of eliciting specific behavioral effects. These results underscore the utility of PFM for informing ES procedures and understanding their large-scale effects on the brain.

### Limitations

Our analyses assume a simplified model of intracranial recordings, which could be expanded on in future studies. We estimated the listening zone of our recording sites using a data-driven procedure (**Supp. Fig. S5**) based on the local correlation of SPES-evoked responses, and from these analyses a 10-mm Gaussian was used in multiple downstream analyses involving Network Dominance. It is possible that these stimulation-evoked responses are larger than what is observed during physiologically coordinated activity, meaning that the “typical” listening zone of implanted electrodes during (non-stimulation evoked) neuronal activity may be smaller than assumed in this study. As a result, multiple networks were often located within this volume surrounding each stimulation site and recording site, increasing the potential complexity of analyses and interpretation of results. To partially account for this, we calculated multiple dominance of targeted network values for each stimulation site. However, we used a simplified winner-takes-all approach to assign each recording site to a single network for interpretability. It is notable that this simplified model still revealed network-level stimulation effects, however, future studies could implement more advanced models, e.g., using electric field modeling (*41*), to better represent the multiple networks likely being recorded from at each recording site.

Additionally, our work focused on the effects of stimulation near functional regions in cortex where the boundary with white matter was well defined and for which there was accurate sampling to the cortical surface, where the fMRI-based networks were defined. We chose to include a minority of MTL and subcortical stimulation sites for earlier analyses that did not assess the distance to the boundary with white matter nor the surface-based networks, but then excluded these MTL and subcortical sites from the rest of the study. Future work will need to assess whether the principals discussed here generalize to MTL and subcortical stimulation, given its clinical relevance.

Finally, a major limitation of this work is that we capitalized on the fact that clinical SPES and HFES procedures typically start at low current intensity and then increase incrementally. This allowed us to assess the influence of current intensity, however, our exploratory analyses of the lowest current intensity applied (1 mA) had comparatively few datapoints. A larger analysis that applies lower current intensity to individually mapped brain networks is needed to confirm the network-specific modulation that our data suggests is achievable through precise network targeting.

## Conclusion

In summary, we demonstrate that precision mapping of large-scale brain networks can be used to target ES to elicit network-level effects. We show that task-free resting-state fMRI-based network mapping based on functional connectivity provides useful information for modulating specific networks, and eliciting specific behavioral effects, with intracranial ES. For this purpose, our results show that individualized estimation of brain networks improves on group-based network maps. Our results suggest that successful targeting requires precise tailoring of the stimulation parameters and location, including keeping current intensity low and applying stimulation to white matter that is proximal to the targeted functional region in the adjacent gray matter. In this context, network-specific modulation may be achieved by limiting the applied current intensity to low values. These results inform future efforts to elicit precise, targeted modulation of specific brain networks and cognitive domains, applicable to both research and clinical settings.

## Materials and Methods

### Participants

Participants were 11 neurosurgical patients aged 23 to 61 years (6 males; mean age = 34.7 years) with intractable epilepsy who were scheduled for invasive monitoring at Northwestern Memorial Hospital (**Supp. Table 1**). Patients provided informed written consent as approved by the Northwestern University Institutional Review Board and were compensated for participation. Participants were invited to attend up to 4 magnetic resonance imaging (MRI) sessions between 3 and 195 days prior to their surgery. Patients with previous resections were excluded, but some of the included patients (P1, P5, P7, P8, and P11) had structural abnormalities observed via MRI by the clinical team. None of the recruited patients had ablations. During invasive monitoring, HFES and SPES was performed as part of routine clinical care, with subsequent research SPES sessions for some patients. In some cases, clinical SPES or HFES caused seizures and these stimulation sites were excluded from all analyses and were not stimulated in research sessions, which focused only on sites approved for stimulation by the clinical team.

### MRI acquisition

Scanning was performed at the Center for Translational Imaging at Northwestern University on a 3T Siemens Prisma. Patients completed up to eight 7-min resting-state runs per session. Patients were asked to fixate their gaze on a cross at the center of the screen. Five patients completed up to four runs of a language localizer task interspersed in each session that were not analyzed here. The position of the screen and cross were adjusted for each session to ensure a comfortable viewing angle to minimize head motion. Multiple runs of multi-echo blood-oxygenation-level-dependent (BOLD) data were collected in each MRI session using a 64-channel head coil with the following parameters: TR = 1,355 ms, TE = 12.80 ms, 32.39 ms, 51.98 ms, 71.57 ms, and 91.16 ms, flip angle = 64°, voxel size = 2.4 mm, FOV = 216 mm x 216 mm, slice thickness = 2.4 mm, multiband slice acceleration factor = 6. After the first resting-state run of each session, a high-resolution T1-weighted magnetization-prepared rapid acquisition gradient echo was acquired (TR = 2,100 ms, TE = 2.9 ms, FOV = 256 mm, flip angle = 8°, slice thickness = 1 mm, 176 sagittal slices parallel to the AC-PC line).

### MRI quality control

Resting-state BOLD runs with a maximum absolute motion (maxAbs) > 2 mm or a maximum framewise displacement (maxFD) > 0.4 mm were automatically excluded. Any runs with maxAbs > 1 mm or maxFD > 0.2 mm were visually examined and the whole run was excluded if motion was clearly visible. **Supp. Fig. S3A** shows summary fMRI data quality metrics for each participant, demonstrating that patients were able to provide repeated runs of fMRI data with low head motion despite the repeated sessions. A total of between 35 (35) and 189 (147) minutes of resting-state fMRI data were collected (retained) per participant (**Supp. Table 1**). This resulted in a total of 19.3 (15.1) hours of resting-state fMRI data collected (retained), including up to 3.2 (2.5) hours per participant. For patients with multiple T1 images, we estimated the patient’s gray-white matter boundary and pial surface using each image with FreeSurfer (*70*). We then visually inspected each estimate and selected the T1 image with the least boundary/surface estimation errors. Any remaining errors were manually corrected by trained assessors (L.S., A.M.H.).

### MRI data preprocessing

Each patient’s data were processed separately using a custom preprocessing pipeline (“iProc”) described previously in (*60*) and adapted for multi-echo BOLD data in (*62*). This pipeline minimizes interpolations and spatial blurring to preserve anatomical details within individuals. For the two most recent patients (P6, P9), noise reduction with distribution corrected principal component analysis (NORDIC; *93*) was applied to the raw functional data. For all patients, the first nine volumes (approximately 12 seconds) were removed from each run to account for T1 attenuation effects. A mean BOLD template was generated for each individual by averaging aligned data from all the included runs. Functional data were then registered via this mean BOLD template to a 1.0-mm isotropic resolution T1 image that serves as a native space template for each individual. This registration included within-run motion correction, across-run and across-session alignment to the mean BOLD, and alignment to the T1 image, applied as a single interpolation step to minimize blurring.

Brain extraction was performed on the T1 using FSL’s Brain Extraction Tool (BET, FSL v6.0.3). Deep white matter and ventricle masks were projected from MNI space to the native space template to calculate timeseries for white matter, cerebrospinal fluid, and whole-brain signal. These timeseries along with 6 head motion parameters and all temporal derivatives were regressed out of the data using 3dTproject (AFNI version 2016.09.04.1341; *94*). Data were then bandpass filtered at 0.01-0.10 Hz and projected to the fsaverage6 cortical surface (40,962 vertices per hemisphere, *95*). In this step, volumetric data points were propagated to a common spherical coordinate system via sampling from the middle of the cortical ribbon using trilinear interpolation. Surface-projected data were then smoothed along the surface using a 2-mm full-width at half-maximum (FWHM) Gaussian kernel.

After preprocessing, a temporal signal to noise ratio (tSNR) map was created for each participant to examine data quality. For each vertex on the cortical surface, the mean of the preprocessed timeseries was divided by the standard deviation. These tSNR maps were then averaged across runs (**Supp. Fig. S1**).

### Network estimation

The preprocessed data were used to estimate networks by FC using a multi-session hierarchical Bayesian model (MS-HBM, (*65*); **Supp. Fig. S2 & S6**). MS-HBM provides individual-specific network estimates by integrating priors from multiple levels (e.g., group, cross-individual and cross-run variation) to stabilize network estimates. For each patient, we generated clustering solutions at different *k* values (i.e., number of networks) from 14 to 20. Mirroring our prior studies, we searched for the lowest *k* value in which the following previously established networks could be identified (*36*, *61*, *63*, *64*, *67*): default network A (DN-A) and B (DN-B), language (LANG), frontoparietal control network A (FPN-A) and B (FPN-B), salience network (SAL), cingulo-opercular network A (CON-A) and B (CON-B), dorsal attention network A (dATN-A) and B (dATN-B), premotor-posterior parietal rostral network (PM-PPr), somatomotor network A (SMOT-A) and B (SMOT-B), auditory network (AUD), and visual-central (VIS-C) and visual-peripheral network (VIS-P). Network names were chosen based on (*67*).

Across individuals, lower *k* levels (*k*=14 & 15) did not separate primary visual, auditory, and sensorimotor regions from one another. At higher *k* levels (k=17–20) LANG, DN-A, and/or DN-B became over-split, either splitting hemispherically or otherwise losing key network regions that diverged from their expected organization (e.g., *36*, *67*). Thus, we selected a *k* value of 16 to preserve these three networks while still allowing us to identify as many networks as possible from the list above (**Supp. Fig. S2**). At this *k* value, we were not able to separate FPN-A from FPN-B, nor VIS-P from VIS-C, so note that further substructure may be present in these estimates. Further, because subjects showed inconsistencies in the spatial arrangement of SMOT-A and SMOT-B (e.g., compare participant P4 with P6 in **Supp. Fig. S2**), these two networks were combined into a single “SMOT” network that covered the somatomotor strip. Of the 16 networks, 2 were overlapping regions of low tSNR in the anterior medial temporal lobe (MTL) and/or ventromedial prefrontal cortex (vmPFC) and were likely capturing zones of signal dropout (**Supp. Fig. S1**). These were labeled as noise and were ignored.

Lastly, we observed speckling (i.e., scattered, small clusters of vertices) in our network estimates (**Supp. Fig. S2**). These were particularly evident in lower tSNR regions such as the temporal pole and MTL (**Supp. Fig. S1**) and adversely affected some of our derived metrics, particularly of distance to nearest network region. We therefore ignored network clusters comprised of 10 vertices (∼10 mm^2^) or less, which is smaller than the approximate effective resolution of our data (2.4-mm isotropic voxels, smoothed with a 2 mm FWHM Gaussian kernel). This allowed us to remove the majority of speckles (**Supp. Fig. S3B**). Final network maps after combining the two SMOT networks and then de-speckling are shown in **Supp. Fig. S6.**

Although patients generally provided good tSNR (**Supp. Fig. S1**), it is possible that lower tSNR may have accounted for this speckling and the need to combine the SMOT networks in this population. Patients with the lowest tSNR (P2, P7, P8, P10) also scored worse on motion metrics for each run (**Supp. Fig. S3A**). Some patients (P2, P3) only provided 35 mins of good data (**Supp. Table 1**), which is less data than typically desirable for PFM (≥ 60 minutes).

### Electrode implantation and localization

Depth and/or subdural electrodes were temporarily implanted intracranially in the context of the recommended invasive diagnostic investigation to capture seizure activity (details shown in **Supp. Table 2**). Participants were implanted with left-hemisphere (n = 7), right-hemisphere (n = 2) or bilateral (n = 2) electrodes.

A post-operative CT image was used for localizing electrode positions using iELVis (*96*). The CT image was linearly registered to the pre-operative T1 image (see *MRI acquisition*). Coordinates for each contact’s centroid were determined using BioImage Suite (*97*). For subdural electrodes, contact coordinates were corrected for post-implantation brain shift (*98*). Coordinates were then used to localize contacts with respect to brain anatomy and functionally defined networks.

We took steps to minimize or otherwise account for the errors that can accumulate during the localization process. While using a post-operative CT image registered to a pre-operative T1 image to localize electrodes is the state-of-the-art (*99*), imperfect registration can result in contact coordinates being misaligned with the patient’s anatomy. Thus, results of the registration procedure were carefully checked for each patient. Additional misalignment can be caused by the brain shift that occurs during surgery, which can be more than a centimeter for grid and strip electrodes (*100–103*). Grid and strip localization errors due to brain shift was corrected for, but note this may still leave residual errors (*98*). Additionally, this procedure does not correct the coordinates of depth electrodes, which may also lead to brain shift. Lastly, a loss of precision occurs when translating contact coordinates into native (voxel) space. To address this last issue and limit summation of errors, we initially used a 0.1-mm isotropic voxel space to identify coordinates for each electrode contact. To acknowledge that the resulting contact location is approximate and take into consideration the brain regions surrounding the contact coordinates, we generated regions of interest (ROIs) surrounding each electrode contact. These were made in a 0.5-mm isotropic voxel space due to computational constraints (i.e., coordinates were rounded to the nearest 0.5 mm increment during this process). Bipolar pairs of contact coordinates were then modelled using either spherical or hourglass-shaped ROIs (see *Bipolar montage and contact modeling*). Despite these steps being taken, the locations of implanted electrodes should be seen as approximate, along with the reported metrics.

### Bipolar montage and contact modeling

A bipolar montage was created for each patient. For depth and strip electrodes, bipolar pairs were made between neighboring contacts. For grid electrodes, we first divided the grid into strips along its longest dimension and then created bipolar pairs between neighboring contacts within each longitudinal strip. In some cases, however, clinical bipolar stimulation was delivered between contacts across longitudinal strips. These additional bipolar pairs were incorporated into the montage but were used only for modeling those stimulation sites and were not used as recording sites.

To characterize the anatomical location of each bipolar pair, a 2-mm radius sphere was generated around each contact coordinate contributing to the bipolar pair, producing an hourglass-shaped ROI for each bipolar pair. Note that we used the same sized spheres for both depth contacts and strip/grid contacts which are substantially larger (Depth: 1.32-2 mm wide, Grid/Strip: 4 mm wide) and further spaced apart (Depth: 3.5-5 mm spacing, Grid/Strip: 10 mm spacing). We referenced this hourglass ROI to the patient’s cortical segmentation derived from FreeSurfer (*70*). Voxels in the ROI were labeled by tissue type: white matter, gray matter, ventricles, or subcortex. Bipolar pairs in which one of the spheres was entirely outside of the brain were excluded.

For stimulation sites, each bipolar pair was assigned to the tissue type with which the ROI most overlapped with using winner-takes-all. For recording sites, where our focus was on responses of networks located in gray matter, bipolar pairs that were located entirely in white matter were excluded, and the remaining montage was used to estimate the size of the listening zone of recording sites (**Supp. Fig. S5A-C**; see *Estimation of listening zone for recording sites during SPES*) and for replication of the relationship between SPES-evoked responses with BOLD correlation (**Supp. Fig. S5F;** see *Replication of SPES effects*). Note that this montage retained a minority of recording sites near the MTL and subcortex, which are subject to inaccurate network assignment. All SPES analyses were reran after excluding MTL and subcortical recording sites and the resulting findings remained the same.

After estimation of the listening zone and analyses replicating the SPES-BOLD correlation, we excluded stimulation sites near the subcortical midline and medial temporal areas from the remaining analyses. This was done because the gray-white matter boundary is poorly defined by FreeSurfer (*70*) in these areas, making it difficult to estimate distance to this boundary accurately. Further, in many cases our fMRI-based definition of the networks of interest was performed on the cortical surface, which omits or improperly samples medial temporal lobe structures. To exclude these stimulation sites, gray matter voxels in each site’s hourglass ROI were labeled according to anatomical region. If the stimulation site mainly overlapped with the amygdala, hippocampus, and parahippocampal gyrus cortical segmentation labels from FreeSurfer, it was deemed to be in MTL and excluded from further analysis. Any stimulation sites that previously were assigned to the subcortex were also excluded at this point. Note that this is a limitation of this study which specifically focuses on cortical stimulation.

To assess how the anatomical location of stimulation sites influenced effects, we calculated the distance between the centroid of each bipolar pair and the nearest gray-white matter boundary estimated by FreeSurfer (**Fig. 1B**). Depth relative to the boundary was the distance value, and this was set to negative if the bipolar pair was predominantly in white matter (as defined using the hourglass ROIs) and to positive if predominantly in gray matter. We also calculate the orientation of the bipolar pair with respect to the nearest cortical column (*46*). This procedure involved making a line between the two contact coordinates of the bipolar pair, then estimating the angle subtended between this line and the nearest cortical column.

### Network characteristics of bipolar sites

To contextualize each bipolar site with respect to the functionally mapped networks in the surrounding area, we first calculated the Euclidean distance from each bipolar centroid to each network. To do this, a map of the Euclidean distance from the centroid of each bipolar pair was generated in 0.5-mm isotropic voxel space and projected to the cortical surface.

On the surface, we calculated the nearest distance between the bipolar centroid and the nearest vertex of each FC-derived network (**Fig. 1C**). Thus, for each stimulation site we calculated multiple distances (i.e., one to each network).

Often, a bipolar pair would be near multiple large-scale networks. However, one network would often be more dominant, in the sense that more nearby vertices were assigned to that network than others, or that one network was more closely aligned with the centroid. To capture these scenarios, we derived a metric to represent the “Network Dominance” at each bipolar pair. Dominance was calculated by generating a 10-mm FWHM Gaussian (∼12.5 mm radius; see *Estimation of listening zone for recording sites during SPES)* in 0.5-mm isotropic voxel space centered on the centroid of each bipolar pair and then projecting this to the cortical surface. We calculated the weighted percentage of network-labeled vertices within the Gaussian to derive a Network Dominance value for each network per site (**Fig. 1D**). Note that we use the term “dominance” for simplicity, but that networks covering a minority of vertices at a given site are still discussed in terms of their (low) levels of dominance.

As described above, we characterized stimulation sites in terms of their depth into white matter, orientation, distance to each network, and dominance of each network. To reduce the combinatorial problem of considering multiple networks at each stimulation and recording site, and improve interpretability, we simplified our analysis of evoked responses by assigning each bipolar recording site to the most dominant network in the surrounding region in a winner-takes-all fashion. Recording sites without a major dominant network (i.e., no networks with a Gaussian-weighted dominance above 10%) were excluded from network-related SPES analyses (**Figs. 3-5, Supp. Fig. S7**). Thus, we did not characterize recording sites in terms of their distance to network or depth into white matter. However, by definition, all recording sites used in our analyses were located in or near gray matter (see *Bipolar montage and contact modeling*).

### Single-pulse electrical stimulation (SPES)

SPES was applied using bipolar, biphasic square wave currents (0.50-1 Hz, 300 µs pulse width, 30-120s pulse train). During clinical stimulation, current intensity was increased (in 1-2 mA steps) over consecutive pulse trains and applied using the PE-210AK stimulator (Nihon Kohden, Tokyo, Japan). During research stimulation, clinically approved sites were stimulated using a constant current intensity of 5 mA using the S88 Stimulator (Grass Instruments, Sequim, WA, USA). Stimulation was applied while patients were awake and resting. An epileptologist was present to assess and address induced seizures, after-discharges, or other adverse complications.

During each SPES pulse train, evoked activity was recorded from the remaining contacts. For SPES applied by the research team, the ATLAS Neurophysiology System (Neuralynx, Bozeman, MT, USA) was used with a sampling rate of 20,000 Hz and a bandpass filter of 0.8–500 Hz, and TTL pulses from the Grass Instruments S88 Stimulator were sent to the ATLAS to mark the onset of each stimulation pulse. For SPES applied by the clinical team, the EEG-1200 electroencephalography recorder (Nihon Kohden, Tokyo, Japan) was used with a sampling rate of 1,000-2,000 Hz and a bandpass filter of 0.08-300 Hz, and stimulation artifacts on a channel near the stimulation site were used to mark the time of each stimulation pulse.

Data were downsampled to 1,000 Hz and visually inspected for artifacts. For quality control, we excluded the following from further analysis: 1) noisy or artifact-contaminated channels; 2) channels with regular interictal epileptiform activity, as defined by the clinical team; and 3) time segments that showed artifacts across most/all contacts. The remaining data were epoched into trials covering between 500 ms pre-stimulation and 500-1500 ms post-stimulation. Trials containing absolute values greater than 500 µV, ignoring data from the stimulation artifact period (20 ms pre-stimulation to 20 ms post-stimulation) were visually inspected by trained assessor C.C. and manually rejected if they included interictal discharges.

Data were then bipolar re-referenced (see Bipolar montage and contact modeling). We then Z-scored each re-referenced timeseries using a trial-specific baseline period (500 ms pre-stimulation to 50 ms pre-stimulation). To counteract the variability in signal polarity induced by bipolar re-referencing while still retaining the biphasic nature of stimulation-evoked responses for visualization purposes, a signal flipping procedure was used. If the largest deflection in the trial-averaged response (20-500 ms post-stimulation) was negative, then all trials were flipped so that the trial-averaged peak was positive. Trial-averaged CCEPs were then visualized and manually rejected if they contained prolonged stimulation artifacts (*57*). To focus on recording sites near the gray matter ribbon, bipolar recording sites in which both contacts were entirely in white matter were excluded (see *Bipolar montage and contact modeling*).

Following data exclusion and preprocessing, data were retained from 102 SPES stimulation sites across nine patients (**Supp. Table 1**) over a range of current intensities (1-7 mA), resulting in 17,819 unique responses to stimulation.

### High-frequency electrical stimulation (HFES)

HFES was applied by the clinical team using bipolar, biphasic square wave currents (50 Hz, 300 µs pulse width, 3-10s pulse train) with increasing current intensity (1-2 mA steps) over consecutive pulse trains. During stimulation, the patient was asked to report any subjective feelings in addition to being observed by an epileptologist in the room.

HFES can elicit “gain of function” behavioral effects such as an involuntary movement or sensation, or “loss of function” effects such as slowed reading or muscle weakness (*2*, *3*). Loss of function can only be assessed during active testing (e.g., reading sentences, performing motor tasks). Following standard clinical procedures, stimulation was either applied while the patient was awake and resting or while the patient performed a task relevant to the suspected function of the stimulated brain region. These included tasks assessing language (reading, naming, repeating, counting out loud, reciting the alphabet, singing “Happy Birthday”), comprehension (performing math, following commands, recall), motor control (moving hands, holding arms up, holding gaze forward, holding gaze right), and smell (sniffing). For each pulse train, the current intensity used, task administered, behavioral effects observed, and signs of seizures and/or after discharges were documented by a clinical professional in the room and later coded by author C.C. Any discrepancies or missing information in the documents were clarified post hoc with the clinical team. It is important to note that many of the HFES stimulation sites in our study were in association cortices, where loss of function effects are predominant over gain of function effects (*17*, *104*, *105*). As such, the observations made during this clinical procedure were limited by the selected functional domains that were assessed during stimulation.

Across all current intensities and tasks administered, HFES sites were categorized into two main groups: 1) those eliciting behavioral effects without after discharges, 2) those not eliciting any behavioral effects while reaching the maximum current intensity (depth electrodes: 7mA, subdural electrodes: 15 mA). Any HFES sites that did not meet either of these criteria were deemed inconclusive and excluded. Any HFES sites in which the patient experienced a seizure-related auras were also excluded. This left 221 conclusive HFES sites across 11 patients (**Supp. Table 1**).

HFES sites eliciting behavioral effects were categorized into 12 effect types. Language and comprehension effects were assessed based on deficits caused during a relevant task. The remaining effect types could occur spontaneously or during active testing and were categorized based on the symptoms written down by the clinical team into the following categories: sensorimotor, visual, auditory, version, temperature, breathing, pain, cognition, and emotion. Effect types were sometimes concurrent, such that two or more effects (e.g., sensorimotor and language) could happen during stimulation of the same site. HFES Sites which elicited behavioral effects that spanned multiple effect types will hereby be referred to as “mixed effects”, while those which elicited behavioral effects within a single category will be referred to as “clear effects”. All of the HFES sites causing the full range of effect categories were included in the analysis to assess the effect of the stimulation site’s depth into white matter (**Fig. 6**). Discussed in the next section, a subset of these sites, which caused effects within specific categories, was further assessed in terms of their relation to individual networks (**Fig. 7**, **Supp. Figs S8-S15**).

### Analysis of network-related behavioral effects

To assess if the presence of network regions near the stimulation site increased the incidence of behavioral effects related to that same network, the Euclidean distance from each stimulation site to each surrounding network region (i.e., the “targeted network”) was calculated. This was done only for stimulation sites near the gray matter ribbon (no more than 5 mm into white matter), and networks which we selected based on their relevance to the six effect types (Sensorimotor, Language, Auditory, Visual, Temperature, and Version effects) that were observed as clear effects (i.e. in the absence of other concurrent effects) at more than one stimulation site, to ensure generalizability.

For each stimulation site-network region distance, we categorized the behavioral effect as (i) a clear effect related to the targeted network (“Clear Network *N* Effects”), (ii) a mixed effect related to the targeted network (“Mixed Network *N* Effects”), (iii) a clear or mixed effect not related to the targeted network (“Other Effect(s)”), or (iv) “No Effects”. Language effects were only assessed at HFES sites at which a language task was performed. These stimulation outcomes were then binned by stimulation site-network region distance, and the proportion of outcomes belonging to each behavioral effect category were calculated within each distance bin. We then performed a permutation test to assess the statistical significance of network-related stimulation effects. Behavioral effect category labels were permuted 10,000 times to randomize the relationship between stimulation outcome and distance between the stimulation site and the targeted network region. After each permutation, the data were binned according to distance, to provide a null distribution for the expected proportion of effects for each category within each distance bin based on chance. If the true likelihood of eliciting targeted network effects (i.e., Clear Network *N*

Effects, plus Mixed Network *N* Effects) for a given distance bin was more extreme than 95% of its respective null distribution, it was deemed significant. During this test we only assessed distance bins with at least five observations (i.e. distance bins closer than 55 mm to the targeted network).

### Estimation of listening zone for recording sites during SPES

As groundwork for the analyses of long-range evoked responses of SPES, we estimated the size of the listening zone of recording sites in order to better understand the volume of tissue that each recording site is sampling from. We adapted an analysis from (*106*) which measures the drop off in correlations between task-evoked responses with increasing inter-response distance, applied here to SPES-evoked responses (**Supp. Fig. S5A**). First, to exclude responses near the stimulation site that could be due to volume conduction, we used a data-driven cutoff to exclude local CCEPs. We plotted the maximum amplitude of each trial-averaged CCEP (20-500 ms post-stimulation) against distance to the stimulation site (Euclidean distance between bipolar centroid coordinates from the stimulation site to the response site). **Supp. Fig. S5B** shows that response amplitude began to level off after a distance of ∼20 mm from the stimulation site. Thus, for each stimulation site, recording sites within 20 mm were excluded.

Next, pairwise Pearson’s product moment correlation coefficients were calculated between recording sites for responses 20-500 ms post-stimulation. Following methods from the original paper (*106*) this was done for the bipolar recording sites that were within 30 mm of each other, which were then averaged across trials. A decay function (𝑟 = (1 − 𝐵)^#^) was used to model the relationship between the absolute response site-response site correlation values (𝑟) and inter-response site distance (𝑑) by optimizing the decay factor (𝐵) using a least-squares fit. Using the fitted model, we took the distance at which the correlation value equaled 0.5 to represent the half-width half-maximum of the data, which was doubled to generate a full-width half-maximum (FWHM) value.

This process was repeated to calculate a separate FWHM value for each stimulation site, current intensity, and resulting set of responses. Across response patterns, the recording sites in our study had a median listening zone of 10.71 ± 4.8 mm FWHM (**Supp. Fig. 5C**). This analysis yielded results similar to (*106*) who studied task-evoked responses, where bipolar referenced gray matter bipolar pairs had a mean listening zone of 7.19 ± 1.6 mm FWHM in the lowest frequency band assessed (theta, 4-8 Hz). Therefore, we used a 10-mm FWHM Gaussian as weighting to calculate the Network Dominance surrounding stimulation sites and recording sites.

### Replication of SPES effects

Following (*46*), at each stimulation site, local (within 20 mm of the stimulation site) and distant response amplitudes (trial-averaged maximum amplitude, 20-500 ms post stimulation) from the highest current intensity applied at each stimulation site (5-7 mA) were averaged and then binned by the depth of each stimulation into white matter (**Supp. Fig. S5D**). We also binned recording sites according to the orientation (angle subtended) to cortical columns; **Supp. Fig. S5E**). See *Bipolar montage and contact modeling* for further details.

Following (*13*), we calculated the mean BOLD timeseries for each recording site and stimulation site region (modeling each site as a spherical ROI of 6-mm radius, centered around the virtual bipolar coordinate, generated in 1-mm isotropic voxel space) using preprocessed but unsmoothed BOLD data. Next, Pearson correlation coefficients were calculated for each run between the mean timeseries at each stimulation site and recording site and then averaged across runs. Using SPES data from the highest current intensity applied for each stimulation site (5-7mA), stimulation site-response site pairs were grouped by whether or not they were responsive (trial-averaged maximum amplitude, 20-500 ms post stimulation, Z > 2), excluding response sites within 20 mm of the stimulation site.

## Acknowledgments

This research was supported in part through the computational resources and staff contributions provided for the Quest high performance computing facility at Northwestern University which is jointly supported by the Office of the Provost, the Office for Research, and Northwestern University Information Technology.

We thank the participants and their families for generously volunteering their time to this research. We thank lab members Anna Shinn, Donnisa Edmonds, and Young Hye Kwon, as well as the staff of the Northwestern Memorial Hospital Epilepsy Monitoring Unit for their help with data collection, in particular Mary Margaret Mizera.

This document was written with assistance from AI tools. Specifically, early drafts of the paper were revised for clarity using ChatGPT 4.5. The content was then reviewed and thoroughly edited by the authors. For more information on the nature of AI usage, please contact the corresponding author.

## Author Contributions

Conceptualization: CC, AMH, RMB

Data curation: CC, AMH, ML, LS, RMB

Formal analysis: CC

Funding acquisition: CC, RMB

Investigation: CC, AMH, JJS, ML, LS, VK, JR, SS, RMB

Methodology: CC, AMH, JJS, ELJ, RMB

Project Administration: AMH, LS, RMB

Resources: SML, JEK, JV, VK, JR, SS, RMB

Software: CC, NLA, RMB

Supervision: SS, RMB

Visualization: CC, RMB

Writing – original draft: CC, RMB

Writing – review & editing: CC, AMH, LS, ML, JJS, NLA, JEK, SML, JV, VK, JR, SS, ELJ, CZ, RMB

## Data and materials availability

All data needed to evaluate the conclusions in the paper are present in the paper and/or the Supplementary Materials or will be made available upon publication.

