## Supplemental Material for "Targeting intracranial electrical stimulation to network regions defined within individuals causes network-level effects"

Christopher Cyr *et al*.

**This PDF file includes:**

Supp. Figs. S1 to S15
Supp. Tables 1 and 2

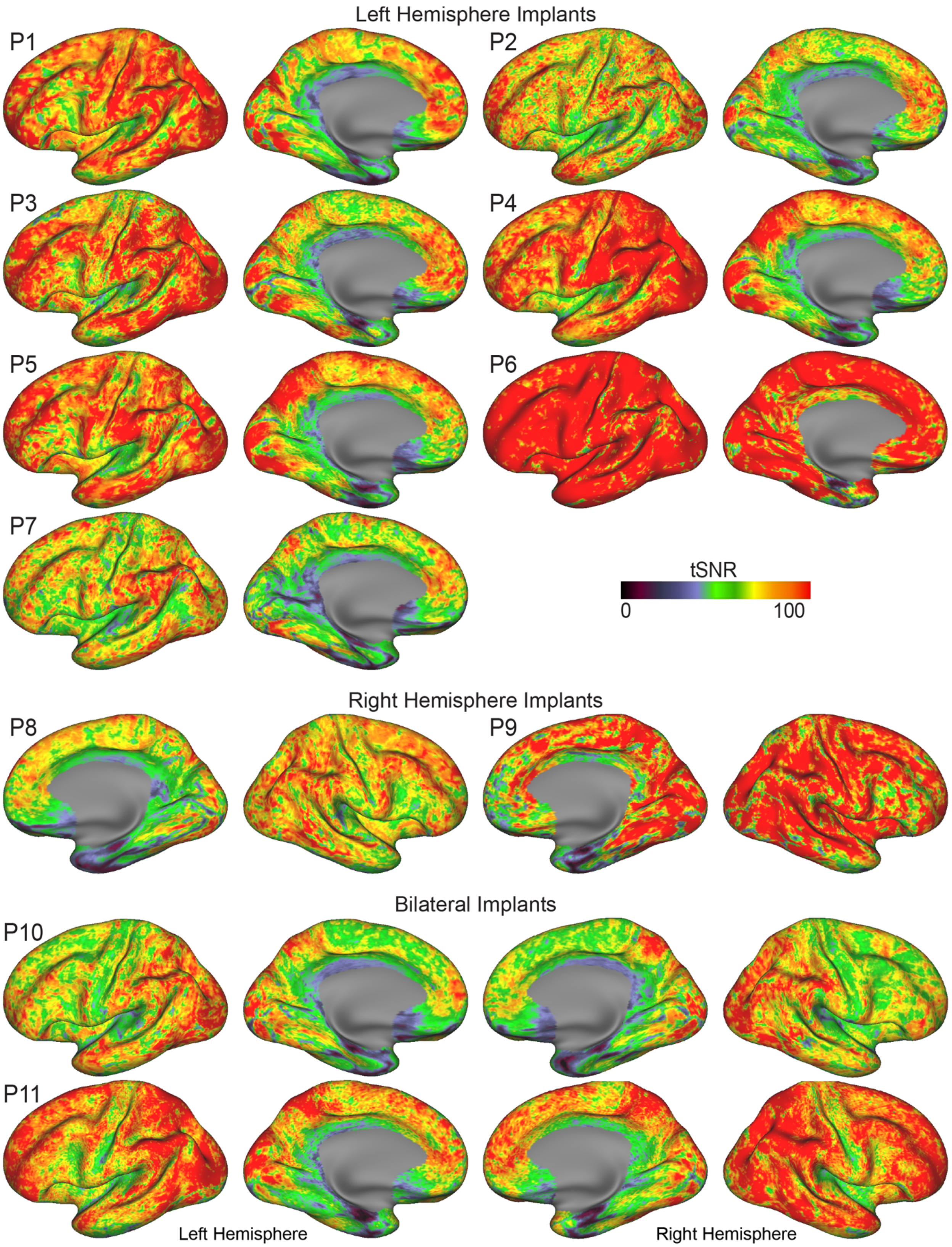


**Supp. Fig. S1. Good temporal signal to noise ratio (tSNR) was achieved throughout the cortical sheet in each participant (P1-P11) using multi-echo fMRI.** The implanted hemisphere(s) is(are) shown for each participant.

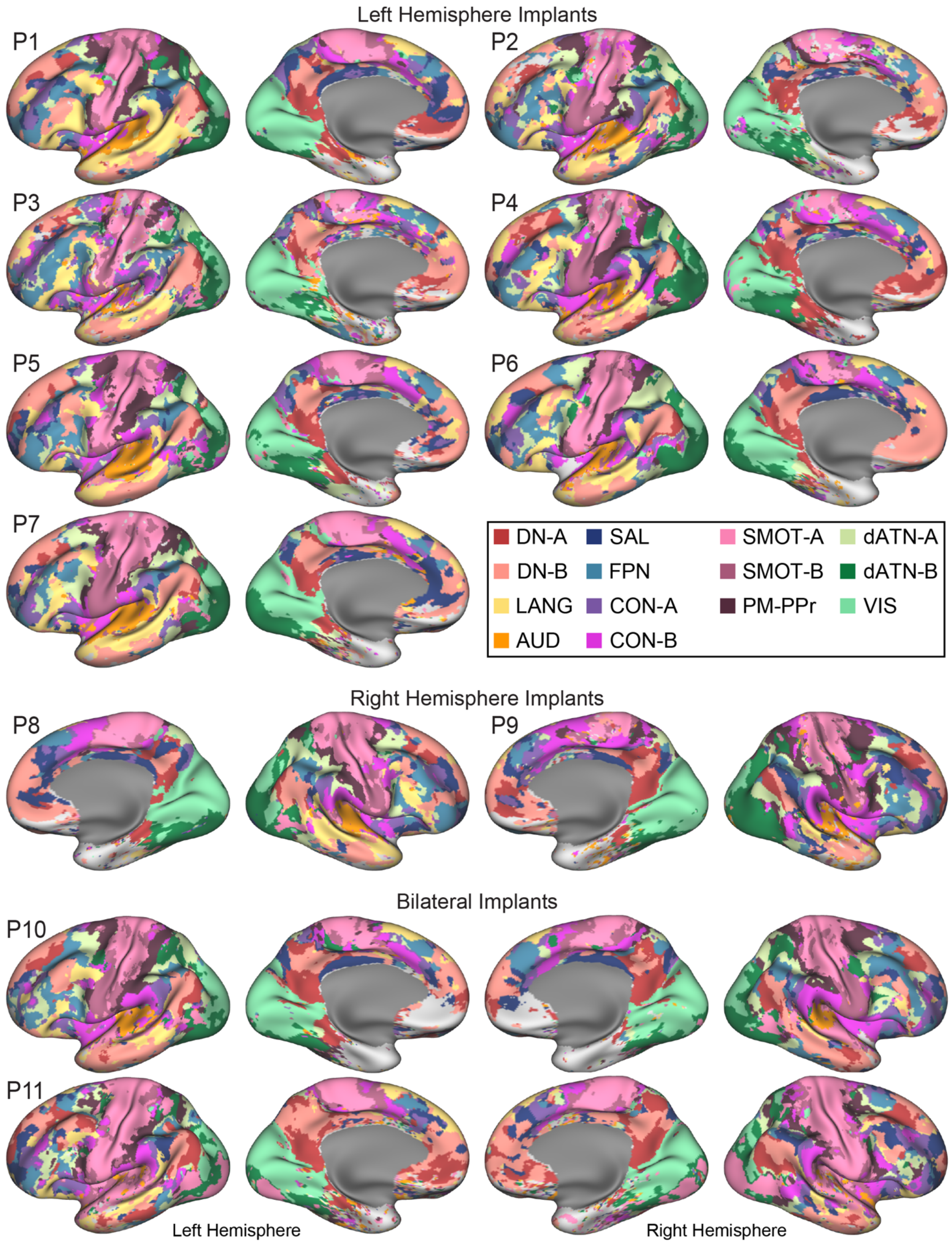


**Supp. Fig. S2: Large-scale networks were estimated within each individual (P1-P11) using precision functional mapping.** Networks were estimated using a data-driven parcellation approach employing a multi-session Bayesian hierarchical model (MS-HBM; *65*). Figure formatted according to Supp. Fig. S1. These estimates later went through two stages of post-hoc correction to combine SMOT-A and SMOT-B networks into one SMOT network due to their inconsistencies across subjects (e.g., compare participant P4 with P6) and to remove small clusters (speckling) from all networks (e.g., see medial temporal pole in P3). Final network estimates are shown in Supp. Fig. S6. DN-A: default network A, DN-B: default network B, LANG: language network, AUD: auditory network, SAL: salience network, FPN: frontoparietal control network, CON-A: cingulo-opercular network A, CON-B: cingulo-opercular network B, SMOT-A: somatomotor network A, SMOT-B: somatomotor network B, PM-PPr: premotor-posterior parietal rostral network, dATN-A: dorsal attention network A, dATN-B: dorsal attention network B, VIS: visual network.

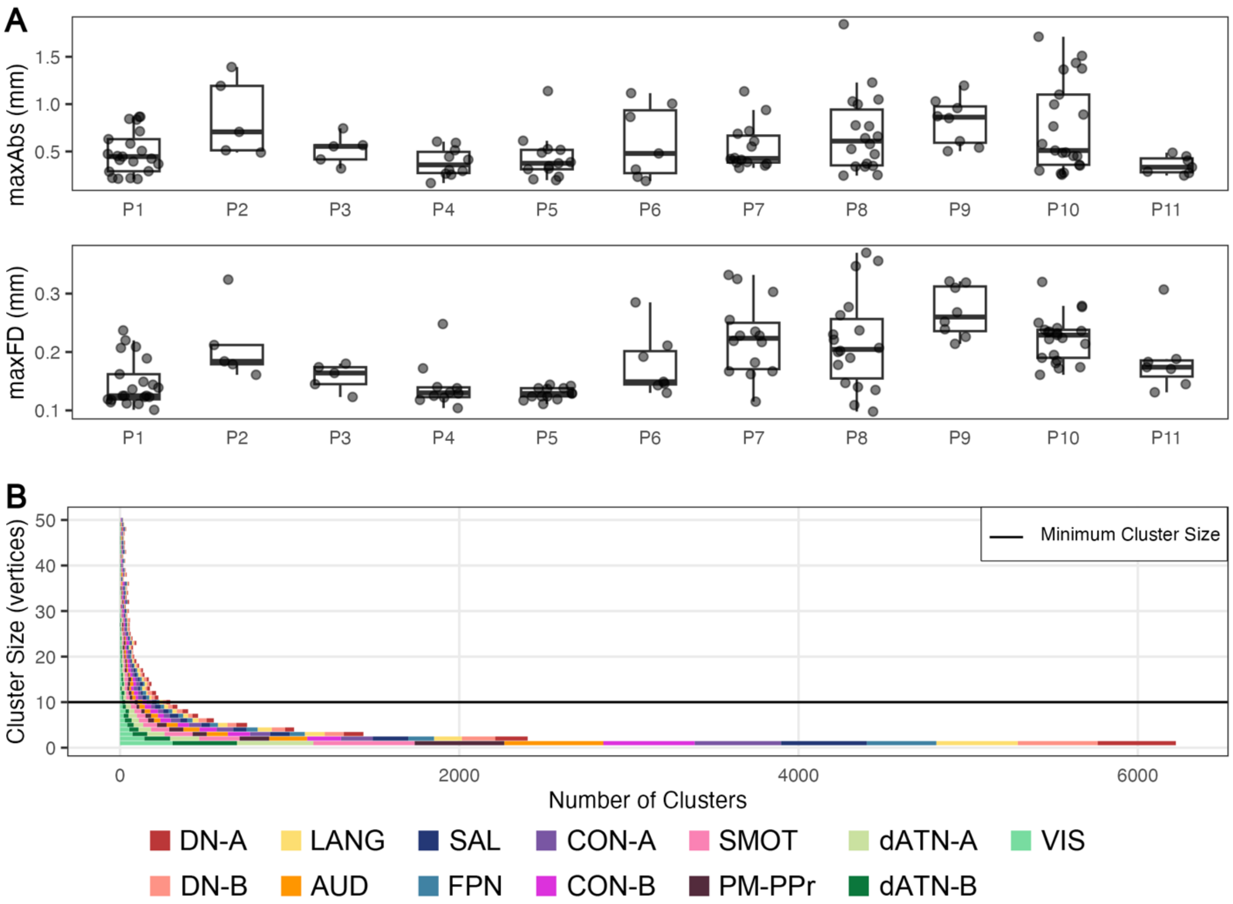


**Supp. Fig. S3: Quality control metrics for fMRI data for each participant and cluster size threshold used for de-speckling.** (A) Maximum absolute displacement (maxAbs; top) and maximum frame-wise displacement (maxFD; bottom) for each resting-state run included in this study, grouped by patient. (B) Distribution of the sizes of individual regions (clusters) across networks, after combining the SMOT networks. To reduce the issue of speckling observed in the MS-HBM results, we ignored clusters composed of 10 vertices or fewer.

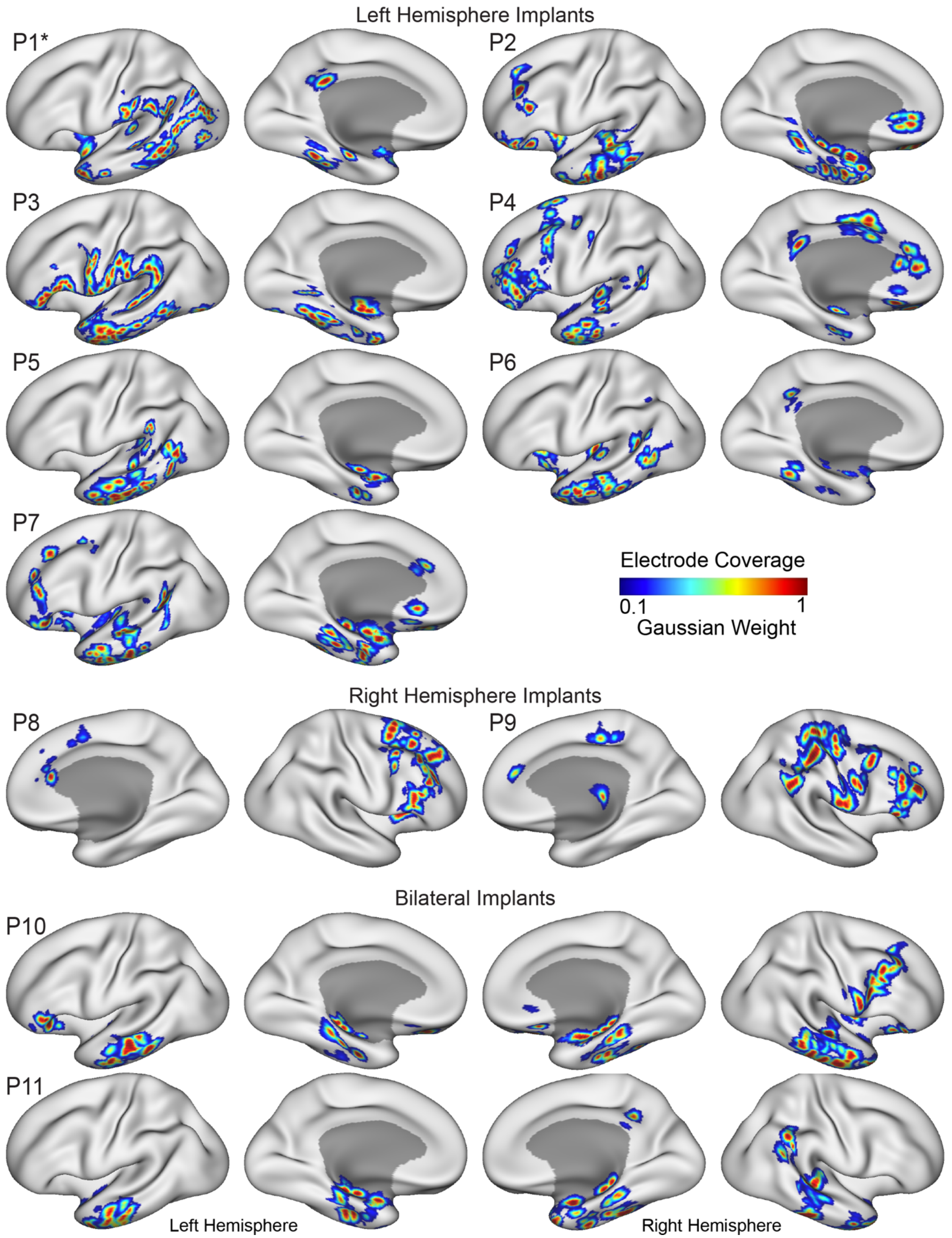


**Supp. Fig S4: Maps showing the estimated locations of each bipolar electrode contact pair for each participant (P1-P11).** Each bipolar pair was modeled using a 10-mm FWHM Gaussian projected to the fsaverage6 cortical surface representation. Coverage varied across individuals but in many cases was broadly distributed. *Patient P1 had two surgeries which are combined here.

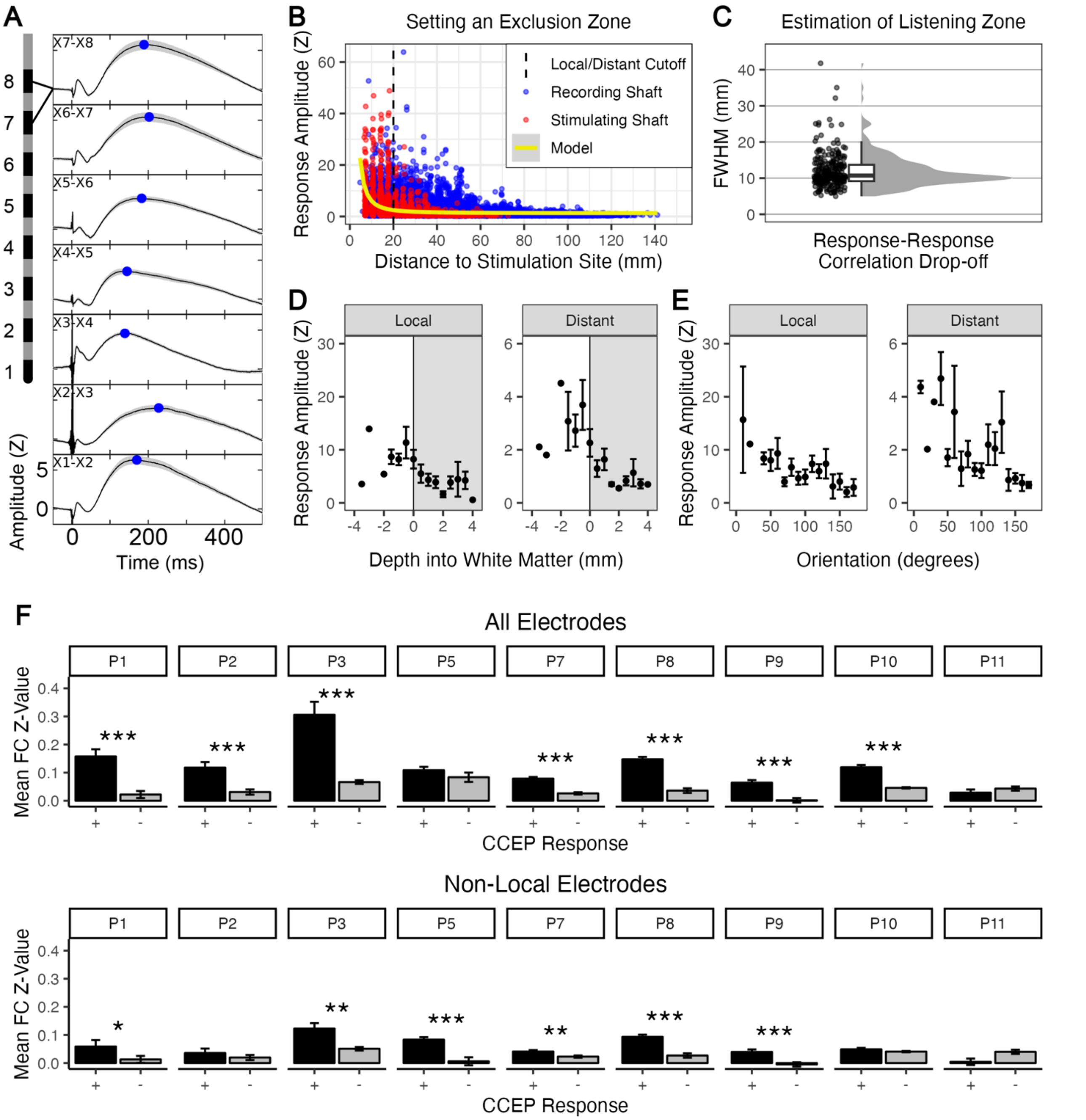


**Supp. Fig. S5: Determining the listening zone of recording sites and replication of expected patterns for single-pulse electrical stimulation (SPES) responses.** (A) An example depth electrode showing correlated responses across neighboring bipolar pairs. Maximum amplitude of the trial-averaged response (20-500 ms post-stimulation) is marked as a blue dot. (B) Plot showing the maximum amplitude of all responses against distance to the stimulation site. A cutoff of 20 mm was used to exclude local responses, which potentially are due to volume conduction. (C) The listening zone of recording sites was estimated using methods adapted from (*106*), with resulting FWHM values across evoked activity patterns resulting from different stimulation sites and current intensities being centered around 10 mm. (D) Replication of findings from (*45*), showing that the amplitude of SPES-evoked responses is higher for white matter stimulation. (E) Failed replication of findings from (*45*) where the orientation of the bipolar stimulation contacts with respect to the gray matter columns did not show an inflection (peak) in local responses near 90 degrees. Note our dataset had fewer stimulation sites than theirs. Local responses are defined as being within 20 mm of the stimulation site. (F) Replication of results from (*13)* showing that stimulation site-recording site pairs with significant CCEPs (“+”), have significantly higher BOLD correlations than those without significant CCEPs ("-"). Asterisks refer to a one-sided t-test between groups within each patient (*$p < 0.05$, **$p < 0.01$, ***$p < 0.001$).

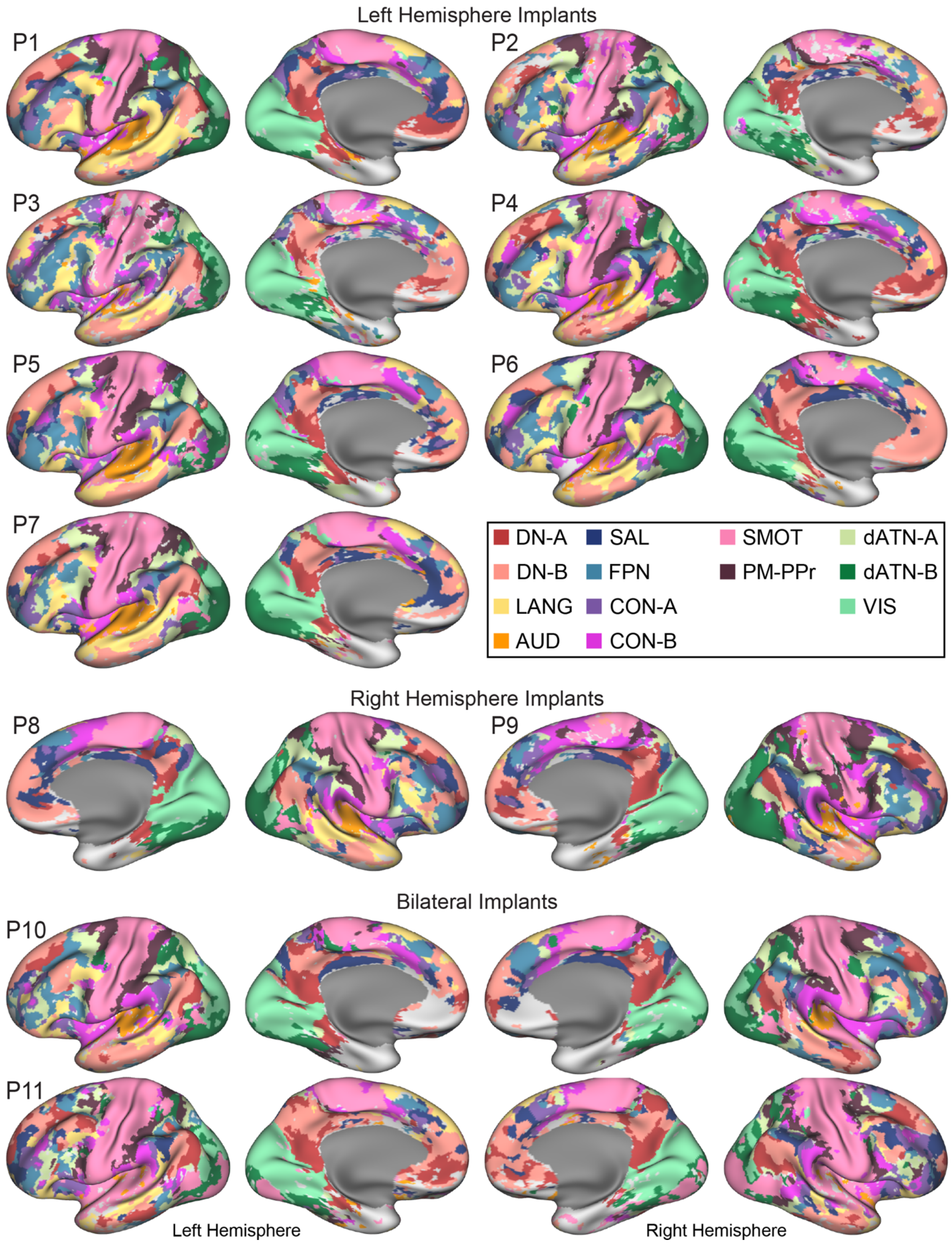


**Supp. Fig. S6: Final fMRI estimates of large-scale networks from all individuals (P1 – P11) after combining networks SMOT-A and SMOT-B and de-speckling.** DN-A: default network A, DN-B: default network B, LANG:

language network, AUD: auditory network, SAL: salience network, FPN: frontoparietal control network, CON-A: cingulo-opercular network A, CON-B: cingulo-opercular network B, SMOT-A: somatomotor network A, SMOT-B: somatomotor network B, PM-PPr: premotor-posterior parietal rostral network, dATN-A: dorsal attention network A, dATN-B: dorsal attention network B, VIS: visual network.

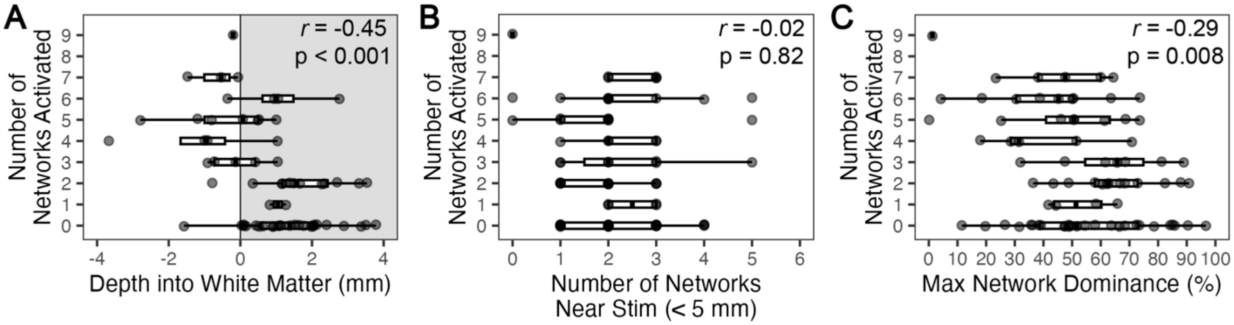


**Supp. Fig. S7: Exploring additional drivers of SPES-evoked responses.** Focusing on instances of 5 mA stimulation, (A) the analysis from Fig. 3F was repeated after removing stimulation sites that were deeper than 15 mm into the brain. This supports that stimulation of peripheral white matter activated more networks than gray matter stimulation, suggesting the white matter depth effect in the full analysis was likely not due to the activation of major (deeper) white matter tracts. (B) To assess whether more networks were activated by stimulation sites near the gray-white matter boundary because these sites were closer to more network regions in the surrounding gray matter (e.g., in both sides of a gyrus), the number of networks with 5 mm of each stimulation site were counted. No relationship was found with the number of networks activated. (C) The number of networks activated did show a relationship with the "Network Dominance" of a single network in the region surrounding the stimulation site. The results of Pearson's correlation tests are shown in the top right corner of each plot.

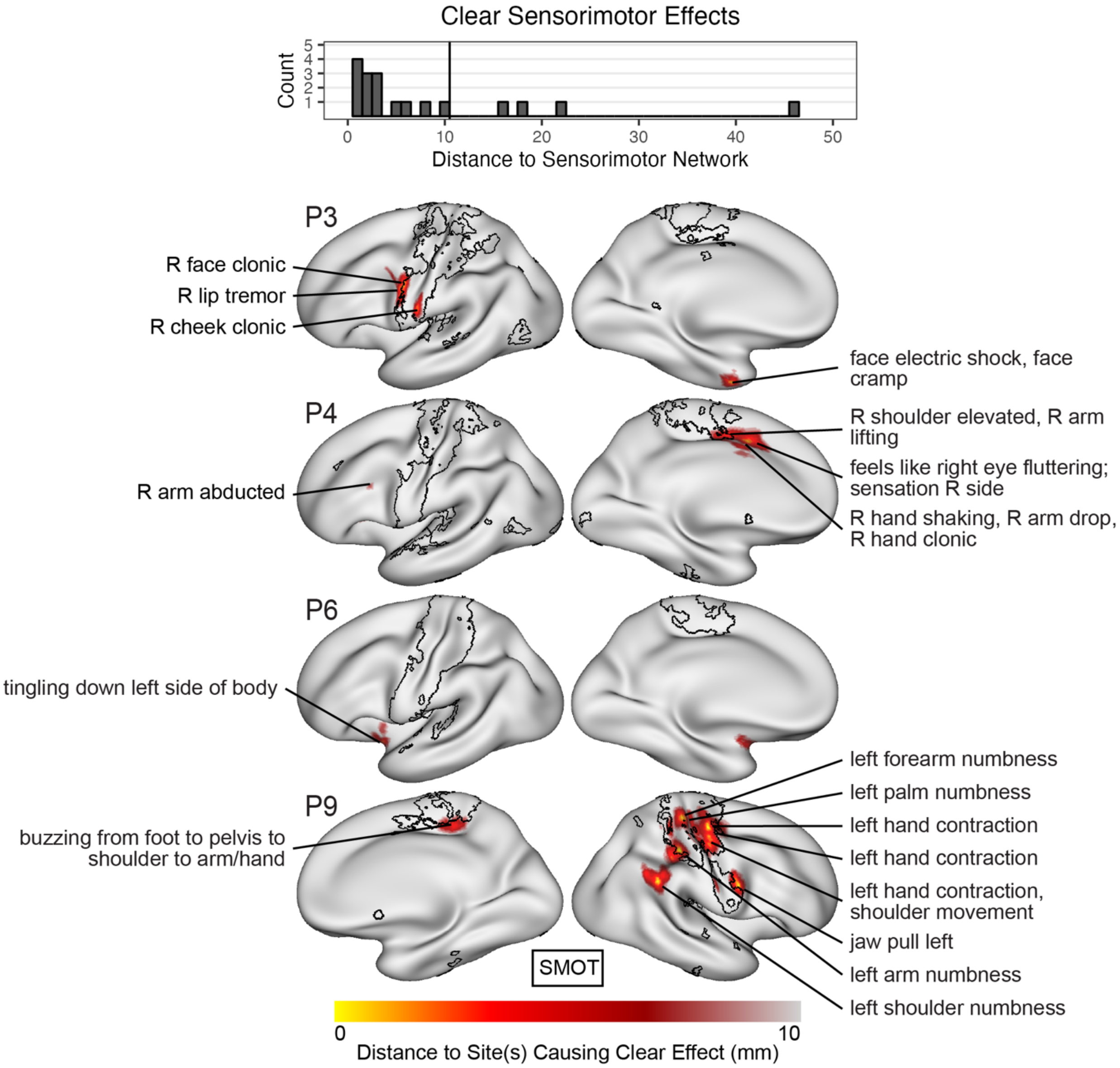


**Supp. Fig. S8: Clear sensorimotor effects occur at stimulation sites near the sensorimotor network.** (Top) Stimulation sites causing clear sensorimotor effects (i.e. only containing sensorimotor components) based on documented symptoms, plotted by Euclidean distance to the sensorimotor network. (Below) Inflated surface views of the cortex showing Euclidean distance to the stimulation site(s) causing effects using the red color bar, with the boundaries of the sensorimotor network (SMOT) in black. Reported symptoms following stimulation of each site are shown. R, Right; L, Left.

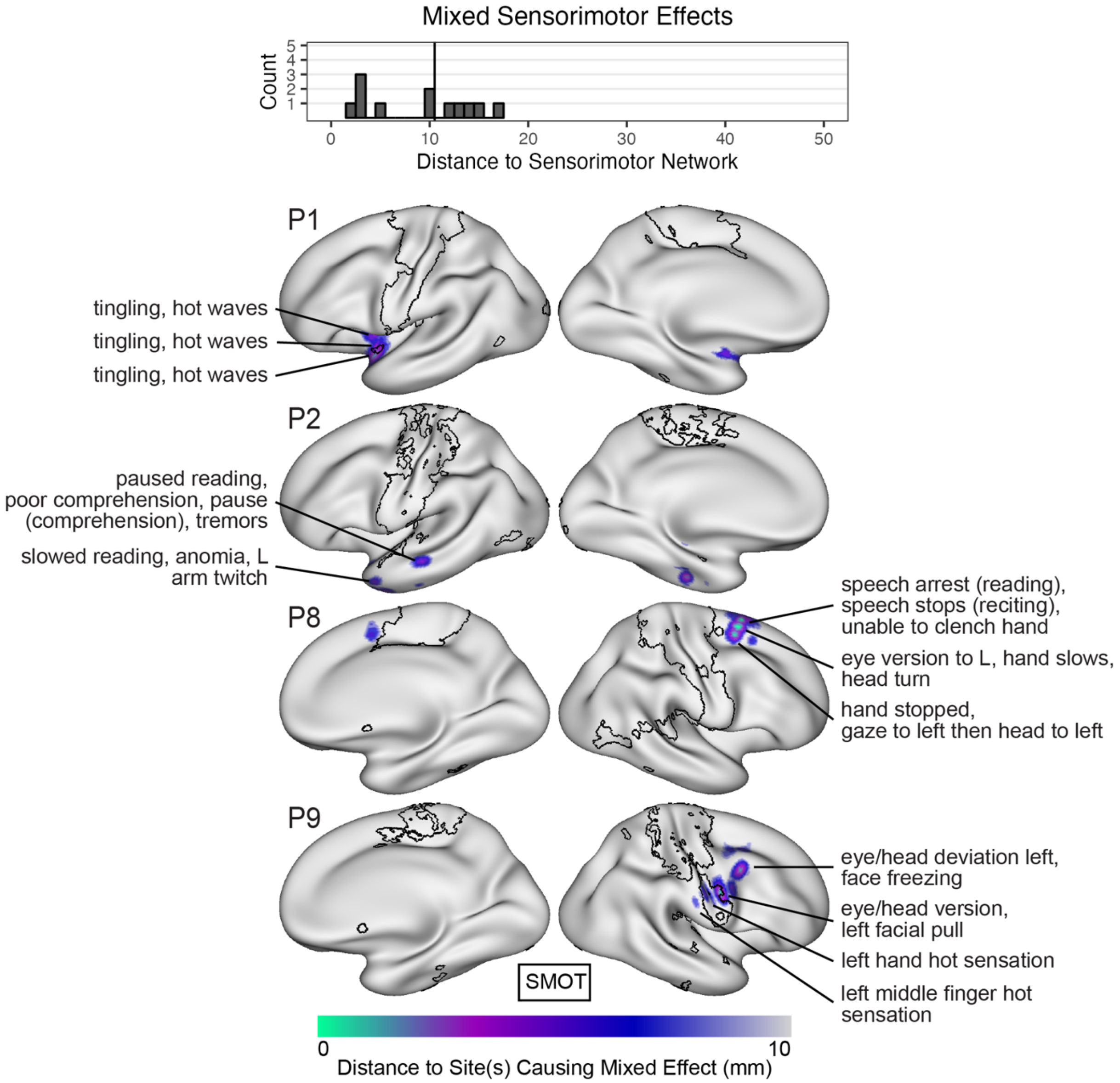


**Supp. Fig. S9: Mixed sensorimotor effects occur at stimulation sites near the sensorimotor network.** (Top) Stimulation sites causing mixed sensorimotor effects (i.e. containing sensorimotor components and non-sensorimotor components) based on documented symptoms, plotted by Euclidean distance to the sensorimotor network. (Below) Inflated surface views of the cortex showing Euclidean distance to the stimulation site(s) causing effects using blue color bar, with the boundaries of the sensorimotor network (SMOT) in black. Reported symptoms following stimulation of each site are shown. R, Right; L, Left.

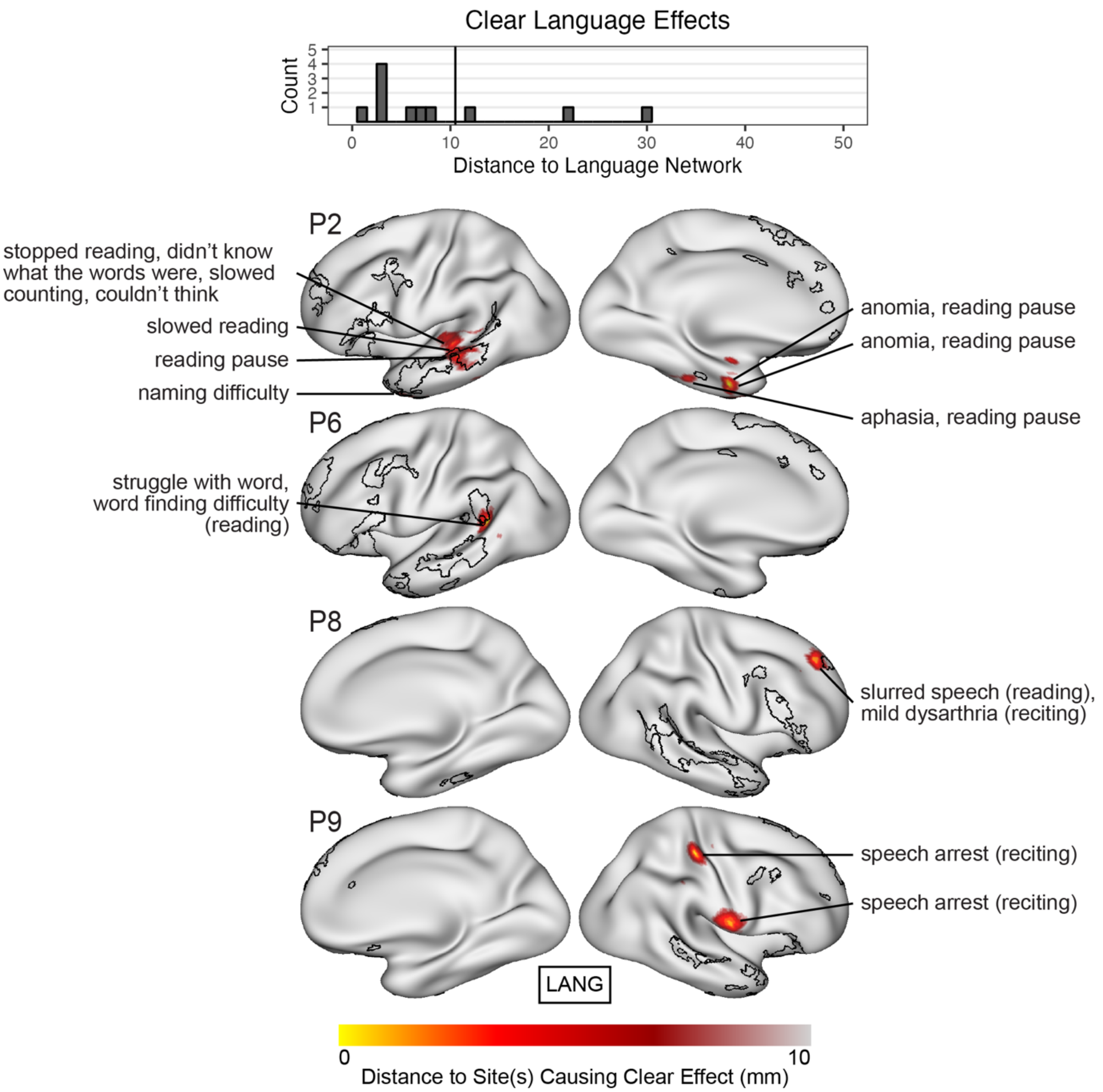


**Supp. Fig. S10: Clear language effects (deficits) occur at stimulation sites near the language network.** (Top) Stimulation sites causing clear language deficits (i.e. only containing language components) during language testing, plotted by Euclidean distance to the language network. (Below) Inflated surface views of the cortex showing Euclidean distance to the stimulation site(s) causing effects using the red color bar, with the boundaries of the language network (LANG) in black. Reported symptoms following stimulation of each site are shown. R, Right; L, Left.

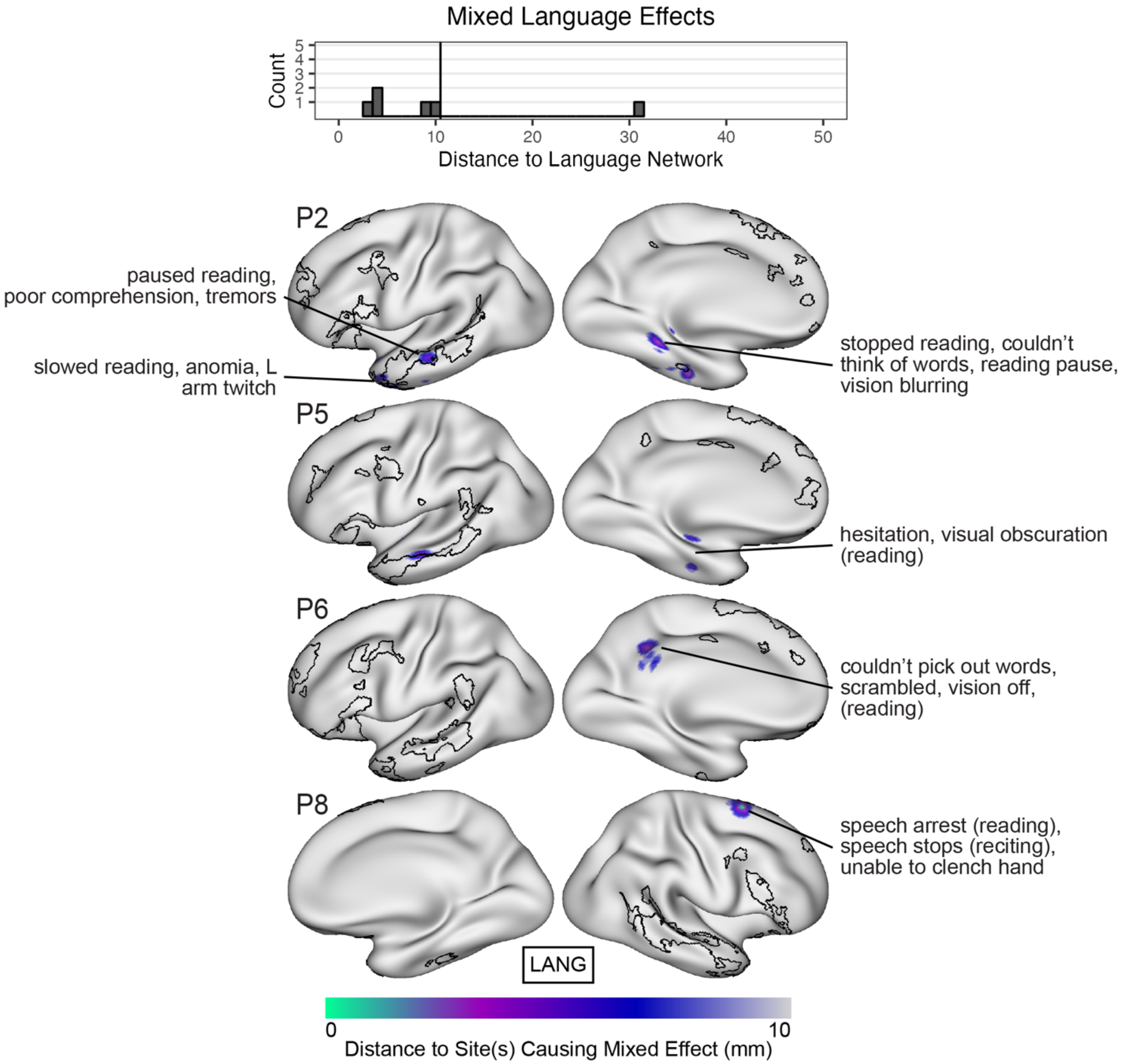


**Supp. Fig. S11: Mixed language effects (deficits) occur at stimulation sites near the language network.** (Top) Stimulation sites causing mixed language deficits (i.e. containing language components and non-language components) during language testing, plotted by Euclidean distance to the language network. (Below) Inflated surface views of the cortex showing Euclidean distance to the stimulation site(s) causing effects using the blue color bar, with the boundaries of the language network (LANG) in black. Reported symptoms following stimulation of each site are shown. R, Right; L, Left.

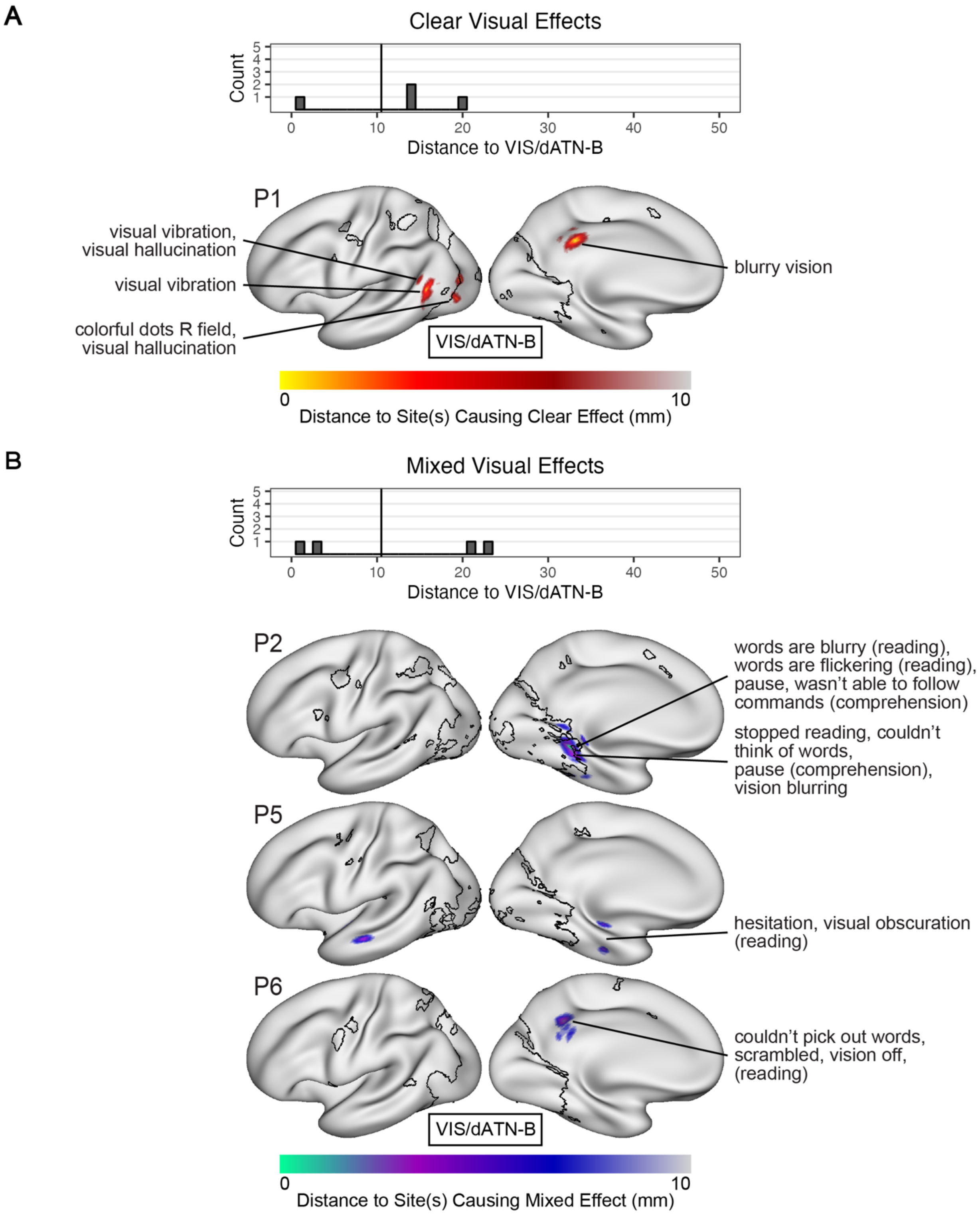


**Supp. Fig. S12: Visual effects occur at stimulation sites near dorsal attention network B and/or the visual network.** Stimulation sites causing (A, Top) clear visual effects (i.e. only containing visual components) and (B, Top) mixed visual effects (i.e. containing visual and non-visual components) based on documented symptoms, plotted by Euclidean distance to the visual and dorsal attention B networks. Inflated surface views of the cortex showing Euclidean distance to the stimulation site(s) causing (A, Below) clear effects using with the red color bar and (B,

Below) mixed effects using the blue color bar, with the boundaries of the visual and dorsal attention B networks (VIS/dATN-B) in black. Reported symptoms following stimulation of each site are shown. R, Right; L, Left.

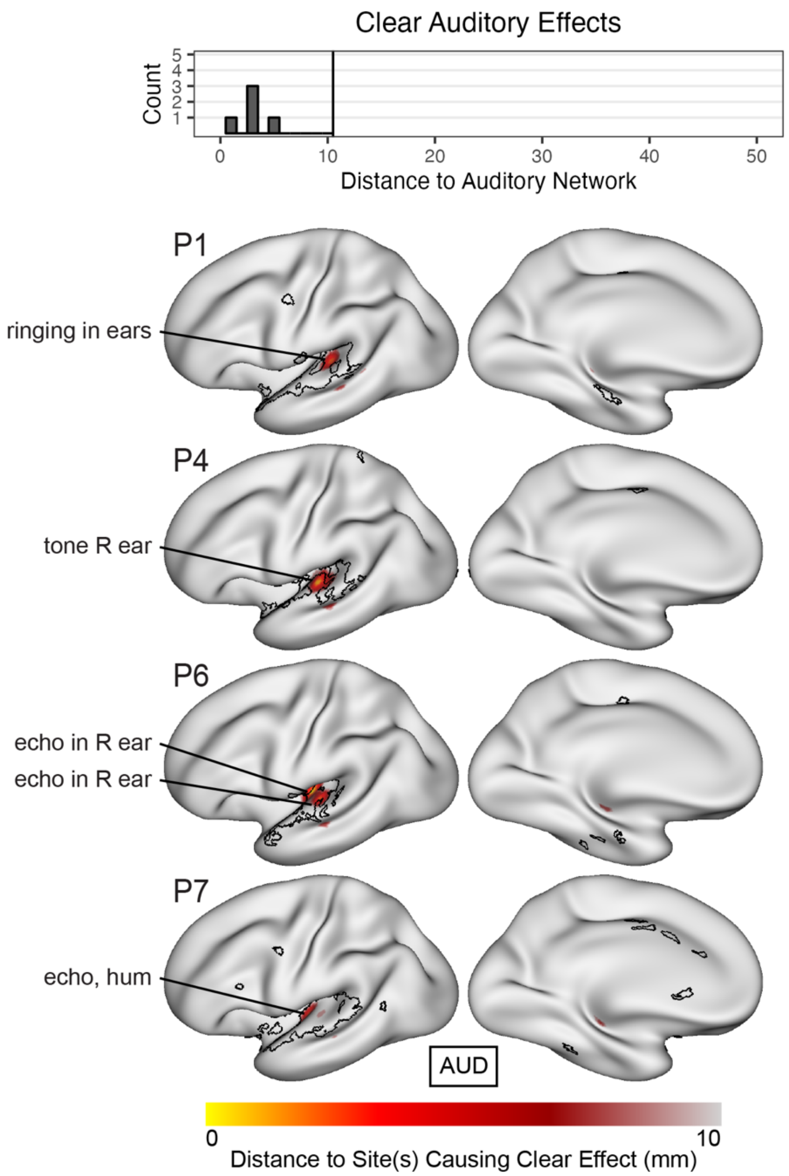


**Supp. Fig. S13: Clear auditory effects occur at stimulation sites near the auditory network.** (Top) Stimulation sites causing clear auditory effects (i.e. only containing auditory components) based on documented symptoms, plotted by Euclidean distance to the auditory network. (Below) Inflated surface views of the cortex showing Euclidean distance to the stimulation site(s) causing effects using the red color bar, with the boundaries of the auditory network (AUD) in black. Reported symptoms following stimulation of each site are shown. R, Right; L, Left.

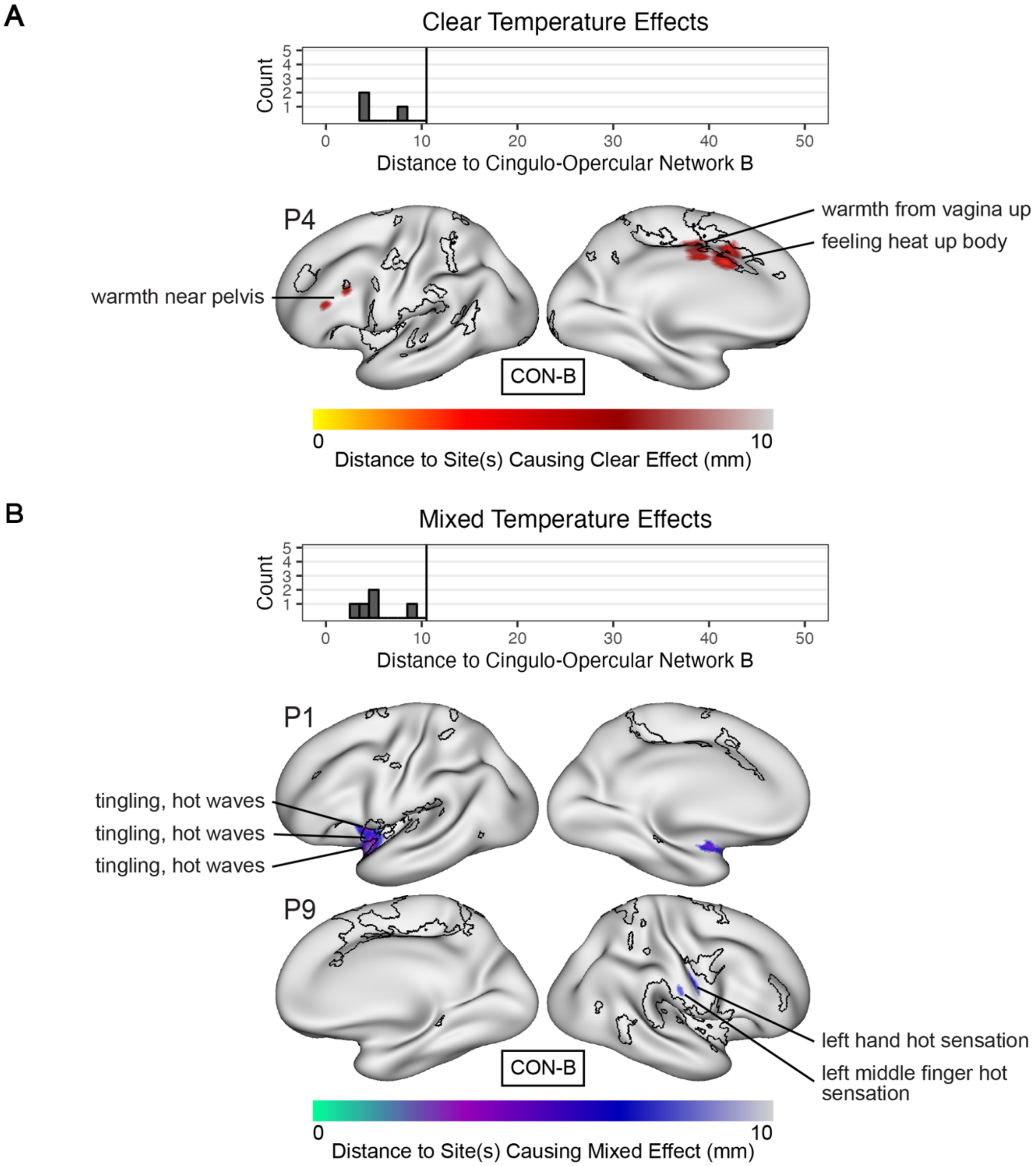


**Supp. Fig. S14: Temperature effects occur at stimulation sites near cingulo-opercular network B.** Stimulation sites causing (A, Top) clear temperature effects (i.e. only containing temperature components) and (B, Top) mixed temperature effects (i.e. containing temperature and non-temperature components) based on documented symptoms, plotted by Euclidean distance to cingulo-opercular network B. Inflated surface views of the cortex showing Euclidean distance to the stimulation site(s) causing (A, Below) clear effects shown with the red color bar and (B, Below) mixed effects shown with the blue color bar, with the boundaries of cingulo-opercular network B (CON-B) in black. Reported symptoms following stimulation of each site are shown. R, Right; L, Left.

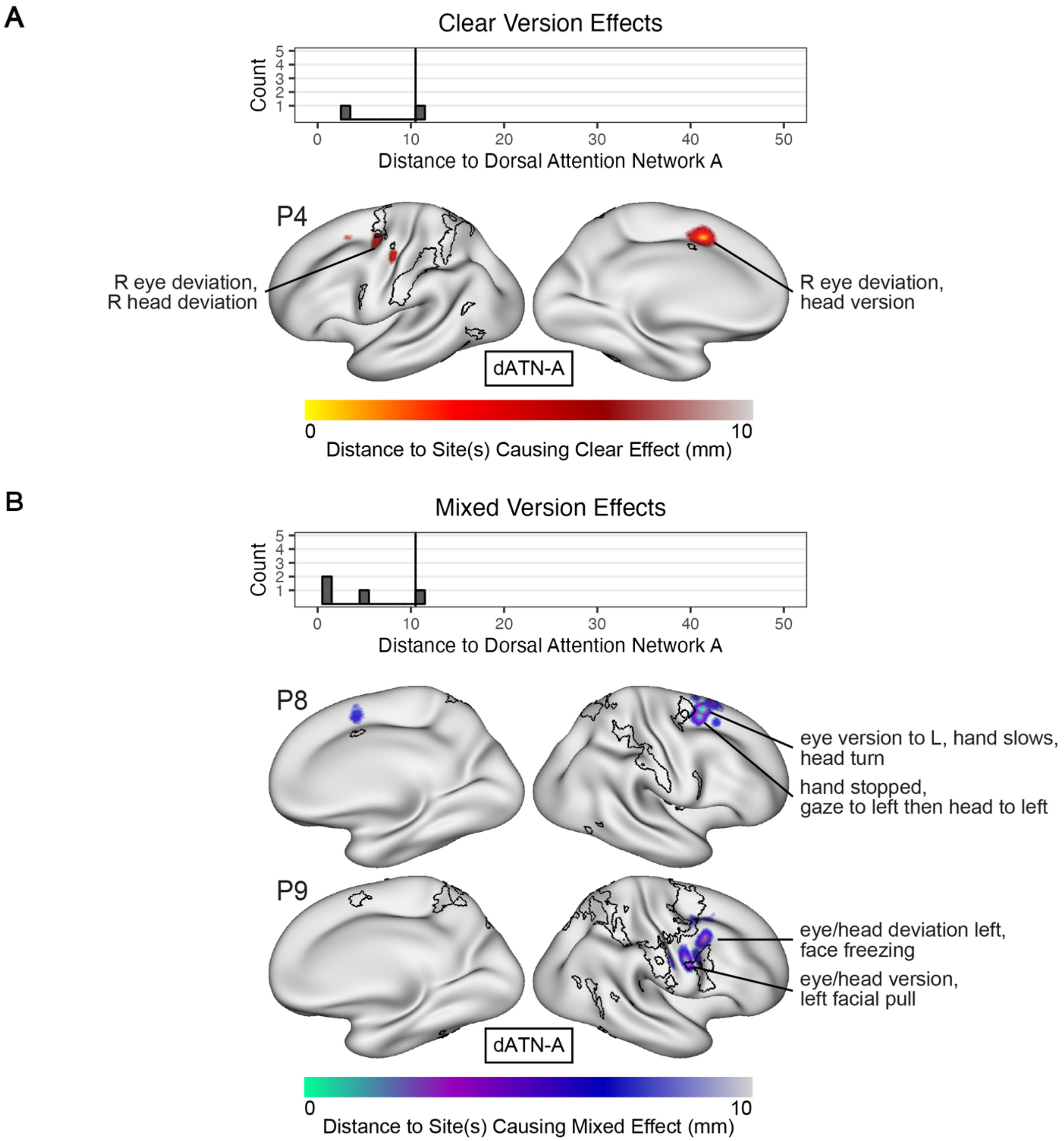


**Supp. Fig. S15: Version effects occur at stimulation sites near dorsal attention network A.** Stimulation sites causing (A, Top) clear version effects (i.e. only containing version components) and (B, Top) mixed version effects (i.e. containing version and non-version components) based on documented symptoms, plotted by Euclidean distance to dorsal attention network A. Inflated surface views of the cortex showing Euclidean distance to the stimulation site(s) causing (A, Below) clear effects shown with the red color bar and (B, Below) mixed effects shown with the blue color bar, with the boundaries of dorsal attention network A (dATN-A) in black. Reported symptoms following stimulation of each site are shown. R, Right; L, Left.

| Patient | P1 | P2 | P3 | P4 | P5 | P6 | P7 | P8 | P9 | P10 | P11 |
|---|---|---|---|---|---|---|---|---|---|---|---|
| | **Demographics** | | | | | | | | | | |
| Age (years) | 23 | 30 | 47 | 27 | 36 | 37 | 33 | 26 | 29 | 33 | 61 |
| Sex | M | M | F | F | M | F | M | M | F | M | F |
| Handedness | R | L | R | R | R | R | R | L | R | R | R |
| | **Clinical Information** | | | | | | | | | | |
| Language Lateralization | LR | L | L | L | L | L | L | L | L | L | L |
| Epileptic Focus | L-TPJ | L-Hipp | L-STG | L-Ins | L-STG L-MTG | L-Amy | L-Hipp L-Amy L-Ins | R-MFG | non-localizable | L-Hipp L-Amy R-Hipp | L-Hipp L-Amy R-Hipp R-Amy |
| MRI Abnormalities | L-Amy Lesion | None | None | None | L-STG L-MTG Dysplasia | None | L-MTL Sclerosis | R-MTL Sclerosis | None | None | L-MTL Sclerosis |
| Epilepsy Duration (years) | 9 | 19 | 18 | 16 | 15 | 7 | 3 | 25 | 16 | 12 | 57 |
| Implanted Hemisphere | L | L | L | L | L | L | L | R | R | LR | LR |
| Electrode Type | Depth Strip Grid | Depth | Depth Strip Grid | Depth | Depth | Depth | Depth | Depth | Depth | Depth | Depth |
| | **Data Summary** | | | | | | | | | | |
| fMRI-Rest (minutes) | 161* (147)* | 56 (35) | 35 (35) | 112 (70) | 91 (91) | 49 (49) | 182 (98) | 168 (126) | 70 (56) | 189 (147) | 49 (49) |
| Electrode Coverage | 198* (65)* | 112 (87) | 88 (76) | 185 (62) | 76 (56) | 159 (27) | 133 (101) | 70 (48) | 112 (64) | 167 (129) | 108 (77) |
| SPES Sites | 17* (13)* | 5 (5) | 13 (12) | 0 (0) | 16 (8) | 0 (0) | 9 (9) | 16 (16) | 11 (11) | 28 (24) | 5 (4) |
| HFES Sites | 71* (26)* | 50 (41) | 33 (7) | 96 (62) | 30 (18) | 54 (27) | 20 (5) | 20 (9) | 21 (15) | 4 (4) | 17 (7) |

**Supp. Table S1. Patient demographics, clinical information, and data summary.** All summary metrics are in terms of the total data collected, with the amount remaining after exclusion and quality control in parentheses. Electrode Coverage, SPES Sites, and HFES Sites are listed in terms of bipolar pairs electrode contacts (one bipolar pair equals one "site"). Electrode Coverage includes stimulation sites as well as recording sites (see Methods-SPES). *Patient P1 had two surgeries which are combined here. L: Left, R: Right, LR: Bilateral; Amy: Amygdala, Hipp: Hippocampus, Ins: Insula, MFG: Middle Frontal Gyrus, MTG: Middle Temporal Gyrus, MTL: Medial Temporal Lobe, STG: Superior Temporal Gyrus, TPJ: Temporal-Parietal Junction.

| Patient | Session | Type | Manufacturer | Number of Electrodes | Dimensions (contacts) | Width (mm) | Spacing (mm) | Diameter/ Exposure (mm) |
|---|---|---|---|---|---|---|---|---|
| P1 | Surgery 1 | Depth | Dixie | 12 | 1x12-15 | 2.00 | 3.5 | 0.80 |
| P1 | Surgery 2 | Depth | Adtech | 2 | 1x10 | 1.32 | 5.0 | 1.12 |
| P1 | Surgery 2 | Strip | Adtech | 1 | 1x6 | 4.00 | 10.0 | 2.30 |
| P1 | Surgery 2 | Grid | Integra | 1 | 4x8 | 4.00 | 10.0 | 2.30 |
| P2 | Surgery 1 | Depth | Dixie | 10 | 1x8-18 | 2.00 | 3.5 | 0.80 |
| P3 | Surgery 1 | Strip | Adtech | 2 | 1x4-6 | 4.00 | 10.0 | 2.30 |
| P3 | Surgery 1 | Grid | Adtech | 3 | 2x6,4x8 | 4.00 | 10.0 | 2.30 |
| P3 | Surgery 1 | Depth | Adtech | 2 | 1x8 | 1.32 | 5.0 | 1.12 |
| P4 | Surgery 1 | Depth | Dixie | 16 | 1x10-15 | 2.00 | 3.5 | 0.80 |
| P5 | Surgery 1 | Depth | Dixie | 8 | 1x10-12 | 2.00 | 3.5 | 0.80 |
| P6 | Surgery 1 | Depth | Dixie | 13 | 1x8-18 | 2.00 | 3.5 | 0.80 |
| P7 | Surgery 1 | Depth | Dixie | 12 | 1x8-15 | 2.00 | 3.5 | 0.80 |
| P8 | Surgery 1 | Depth | Dixie | 8 | 1x8-15 | 2.00 | 3.5 | 0.80 |
| P9 | Surgery 1 | Depth | Dixie | 11 | 1x8-18 | 2.00 | 3.5 | 0.80 |
| P10 | Surgery 1 | Depth | Dixie | 14 | 1x8-18 | 2.00 | 3.5 | 0.80 |
| P11 | Surgery 1 | Depth | Dixie | 11 | 1x8-15 | 2.00 | 3.5 | 0.80 |

**Supp. Table 2: Electrode hardware information for each participant.**
